# A Camera-Assisted Pathology Microscope to Capture the Lost Data in Clinical Glass Slide Diagnosis

**DOI:** 10.1101/2022.08.31.506042

**Authors:** Kimberly Ashman, Max S. Cooper, Huimin Zhuge, Sharon E. Fox, Jonathan I. Epstein, Carola Wenk, Brian Summa, J. Quincy Brown

## Abstract

Digital pathology, or the practice of acquiring, managing, and interpreting high-resolution digital images from glass pathology slides, holds much promise in precision medicine, potentially transforming diagnosis and prognosis based on computational image biomarkers derived from digital tissue images. However, for all its promise, digital imaging in pathology has not yet become an integral part of the clinical workflow as it has in radiology due to high cost, workflow disruptions, burdensome data sizes and IT requirements, and additional dedicated personnel requirements. Consequently, pathology retains the 150-year-old analog workflow, and the vast majority of slides used in clinical diagnosis are never digitized. Furthermore, there is a missed opportunity to capture the image information and associated data on search processes that led to the clinical diagnosis, which could serve as the foundation for computational clinical decision support. This paper describes an approach for slide digitization during clinical review using a camera attached to a standard brightfield pathology microscope. While a pathologist reviews a glass slide using the eyepiece oculars, the continuously running camera digitizes a complete record of the slide review, resulting in multi-resolution slide images and spatiotemporal saliency maps of the slide review. Unlike other approaches, the pathologist does not stop to review the video stream or monitor the acquisition of video frames but performs the diagnostic review at the microscope using the standard clinical protocol. This hybrid analog-digital approach combines the benefits of digital slide analysis, including annotation, computation, and the ability to confirm the completeness and quality of the glass slide review with the ease of using the microscope for primary diagnosis. Furthermore, a record of the pathologist’s attention during the review, including their search path, magnification level, and dwell times at each location on the slide, is obtained. In the future, this approach could enable the development and application of new and emerging computational decision-support algorithms in real-time to provide feedback to the pathologist, reduce diagnostic errors, and improve disease diagnosis and prognosis.

## 1 Introduction

Pathology is at the foundation of disease diagnosis, prognosis, and precision medicine and can be compared to radiology conceptually in that they both involve the analysis of medical images. However, the fundamental difference is that digital imaging is now the *de facto* clinical workflow in radiology. In contrast, the practice of pathology for clinical diagnosis has been analog for over a century, with pathologists reviewing glass slides directly with their microscopes. More recently, two whole slide imaging (WSI) systems have been FDA-approved to scan microscope slides to create digital images of tissue sections for diagnosis.^1^ This has opened up the possibility of telepathology and using computer-assisted algorithms to enhance diagnosis and prognosis.^2–7^ However, less than 1% of clinical pathology labs have replaced traditional microscopy workflows with WSI systems for primary clinical diagnosis, primarily because slide scanning systems are expensive replacements for an inexpensive and efficient microscope (analog) workflow, and their use represents additional steps and delays outside of the current clinical workflow. As a result, the vast majority of slides are never digitized, and most pathologists and patients, especially in lower resource settings, cannot benefit from emerging computational assists in the clinical workflow that require digital images as inputs, such as the recently FDA-approved Paige Prostate.^8^ As new algorithms emerge, their impact on routine patient care will be limited without digital images on which to apply them.

To address the limitations of conventional slide scanners, systems such as the augmented reality microscope (ARM)^9^ are emerging that project computer-generated information, such as annotations or outputs from AI algorithms, onto a microdisplay viewable through the microscope eyepieces in real-time. While these microscopes connect the digital image analysis to the clinical workflow, the results are restricted to the ocular field of view (FOV) at any given time. Without a digital representation of the whole slide, the algorithm and the pathologist are limited to the context provided in a single FOV, which, even at low magnification, may not contain the entire specimen area or region of interest and does not retain any memory of other areas observed by the pathologist that contributed to the diagnosis. Other competing solutions include slide video mosaicking, which either requires an automated microscope stage^10–17^ or requires trained users to turn their attention away from the microscope eyepiece to ‘monitor’ the image acquisition on a computer to ensure non-blurred images.^18, 19^ Such systems slow the pathologist’s natural slide analysis process and alter their trained diagnostic behaviors. To our knowledge, no existing solution provides the capability to “ambiently”^**?**^ generate whole slide images as a pathologist views the slide through the microscope’s eyepieces.

In this work, we aimed to passively create digital slide images, and associated spatiotemporal attention data, from microscope reviews. We captured video streams acquired while a pathologist manually reviewed glass slides with a traditional brightfield microscope, allowing them to change objective lenses at will and move the stage in any direction and at their own speed. We also aimed to demonstrate the suitability of the captured digital slide images for digital image feedback, computational assist, and archival purposes. An essential aspect of our approach is that images collected during the review are captured across resolution scales and combined into a single multi-resolution slide image (MRSI). Additionally, by ensuring that the camera field of view relative to the eyepiece field of view is maximized, and that the camera frame rate is high enough to prevent motion blur or gaps in the final images, our method allows digital recreations of the glass slide review unobtrusively or without altering native pathologist search behavior. As a beneficial consequence, we can derive detailed spatiotemporal saliency maps that reflect the sequence of events and corresponding image data that led to the diagnosis. These data are unavailable in traditional whole slide imaging protocols. In doing so, this system, which we call the Pathology Computer-Assisted Microscope, or ‘PathCAM,’ will connect digital pathology to the established clinical glass-slide workflow, opening up a data pipeline that will fuel current and future advancements in diagnosis and prognosis (Figure 1).

**Fig 1.**
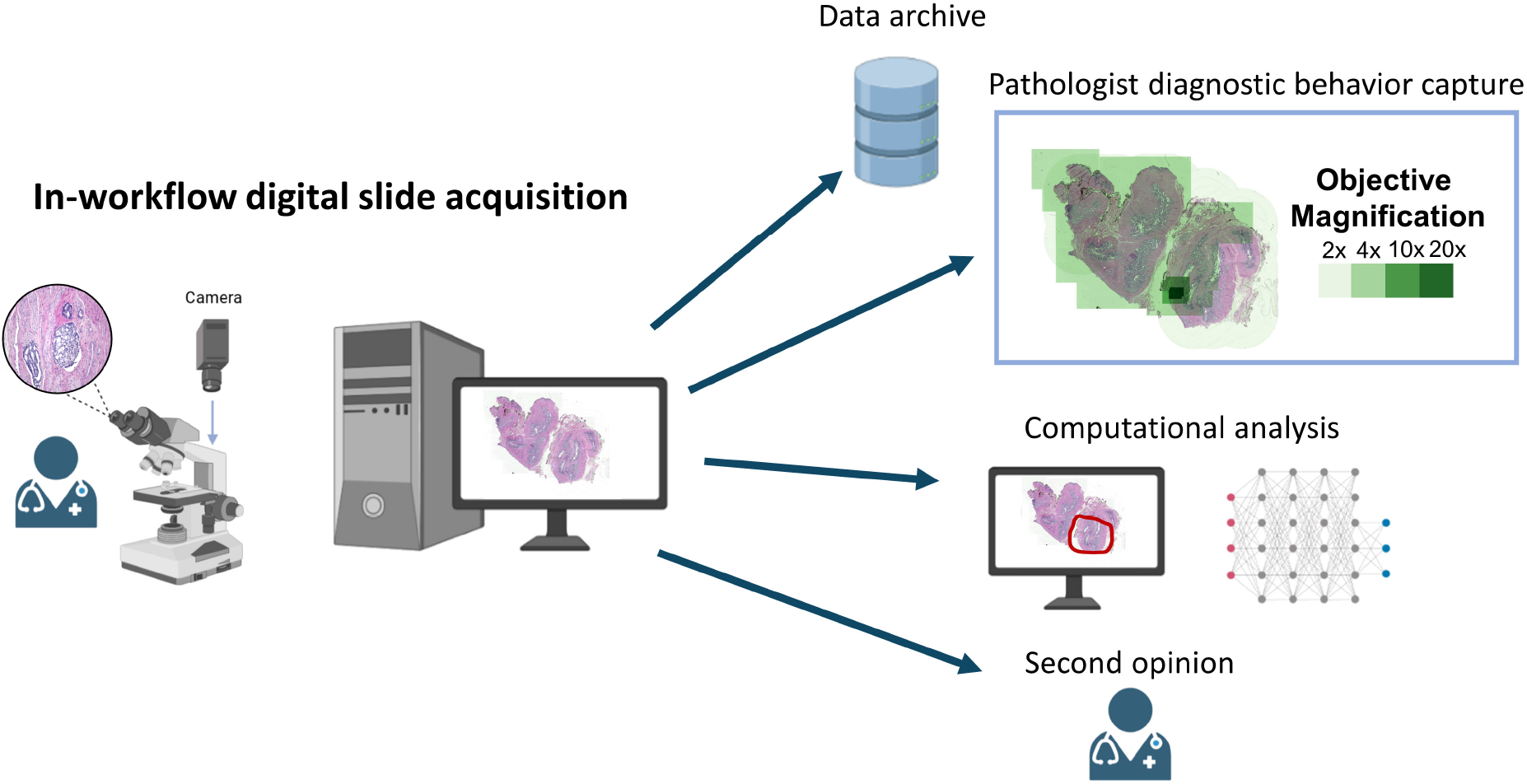
Hybrid analog-digital pathology workflow with a computer assisted microscope. In this re-imagined workflow, we keep the microscope central slide scanning process. Using an onboard camera and video-mosaicking, slides are passively scanned during the pathologist’s microscope analysis. Because there is no change to the pathologist’s behavior, meaning they can move the stage in any direction and change objectives, we capture important information about how the pathologist analyzed the slide. Once the slides are scanned, the captured digital slide images can be used for archival, review, and computational assistance.

## 2 Results and Discussion

### 2.1 PathCAM development and digital slide generation from unconstrained review

Toward our objective of introducing digital capabilities into the existing clinical glass slide workflow, we developed a method of generating mosaics from multiscale pathologist slide reviews without specialized stage hardware. To demonstrate our method, we recorded a pathologist reviewing radical prostatectomy specimens at the microscope, producing slide images using only a commercially available camera and a video-mosaicking routine. We used a microscope with a beamsplitter which enabled simultaneous digitization of the field of view while the pathologist viewed the slide through the eyepieces. We carefully selected the camera sensor such that it was able to digitize nearly the entire FOV seen by the pathologist through the eyepiece oculars, with adequate pixel sampling resolution across all objective lens magnifications used and adequate temporal resolution to capture blur-free images during unconstrained slide movement. Specifically, the camera digitized 86% of the ocular field of view of the objective lenses with a field number (F.N.) of 22 (4x to 40x) and 100% of the FOV of the widefield 2x objective lens with a F.N. of 25, with greater than Nyquist pixel sampling for all of the objective lenses (2x, 4x, 10x, 20x, and 40x). Furthermore, the 31 megapixel camera operated at a maximum frame rate of 26 frames per second (FPS), or a 806 MHz pixel sampling rate, which is sufficient to capture typical slide translation speeds during pathologist review (Supplementary Figure 1).

We captured the video streams while the pathologist reviewed the glass slide through the microscope eyepieces by running the camera continuously. The resulting video streams, which were approximately 76 seconds in duration on average (Figure 2a), were transformed into mosaics through several post-processing steps (described in detail in the Methods section). The average size of the raw captured data was 186 GB per slide recording, but this was reduced to 2 GB (uncompressed) on average per recording through sub-sampling and compositing the frames into mosaics (Figure 2b). Figure 3 shows an example of the mosaic generation process, including frame sampling, flat-field correction, and color correction. For each slide recording, the mosaics produced from different objective lenses were composited into a single, final mosaic, which we call the multi-resolution slide image (MRSI). An example of one of the MRSIs generated from a pathologist’s review of a radical prostatectomy specimen is shown in Figure 4 and Supplementary Video 1. Figure 4a, 4d, and 4e display regions of 4x, 10x, and 2x magnification, respectively, when viewed in the HistomicsUI digital slide viewer (https://github.com/DigitalSlideArchive/HistomicsUI), a widely used digital pathology image viewer developed by Kitware. Whereas during the glass slide review, the pathologist only had access to a single eyepiece field of view at any given time, the PathCAM approach resulted in overview images representing a cumulative digital record of all visual information available to the pathologist during the entire review session. Additional generated slide images are also shown in Supplementary Figure 2. In the following sections, we demonstrate some of the features and capabilities of this new hybrid analog-digital slide review approach.

**Fig 2.**
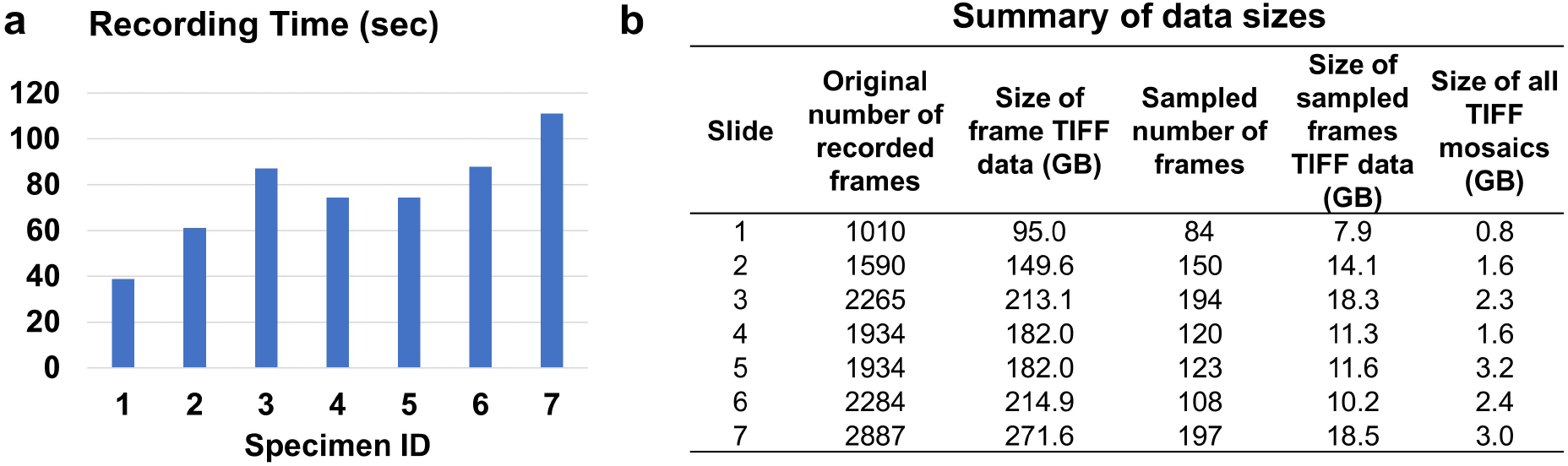
Summary data from digitization of pathologist glass slide reviews. **a**, The recording time of the microscope reviews of each slide. **b**, Summary of data sizes of generated and final frames and mosaics.

**Fig 3.**
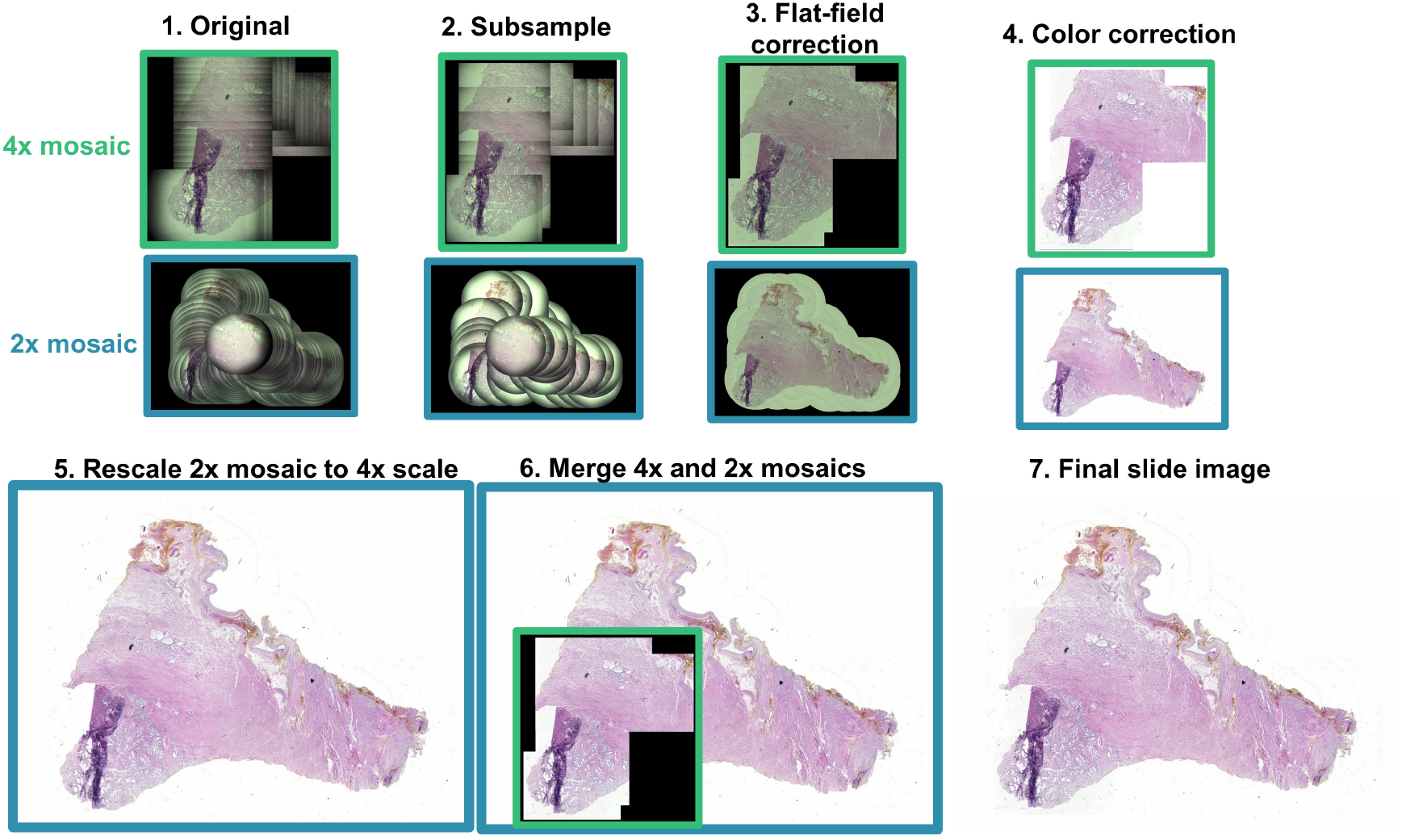
Overview of mosaic creation process. In this example, the mosaics produced from the video recording were subsampled by 10 (b), flat-field (c) and color corrected to create the final mosaics (d). To create a single image from multiscale mosaics, all mosaics are upscaled to match the scale of the largest objective used. For instance, in this example, the 2x mosaic was rescaled by approximately 2 in each dimension and merged with the 4x mosaic to create the final slide image without loss of resolution.

**Fig 4.**
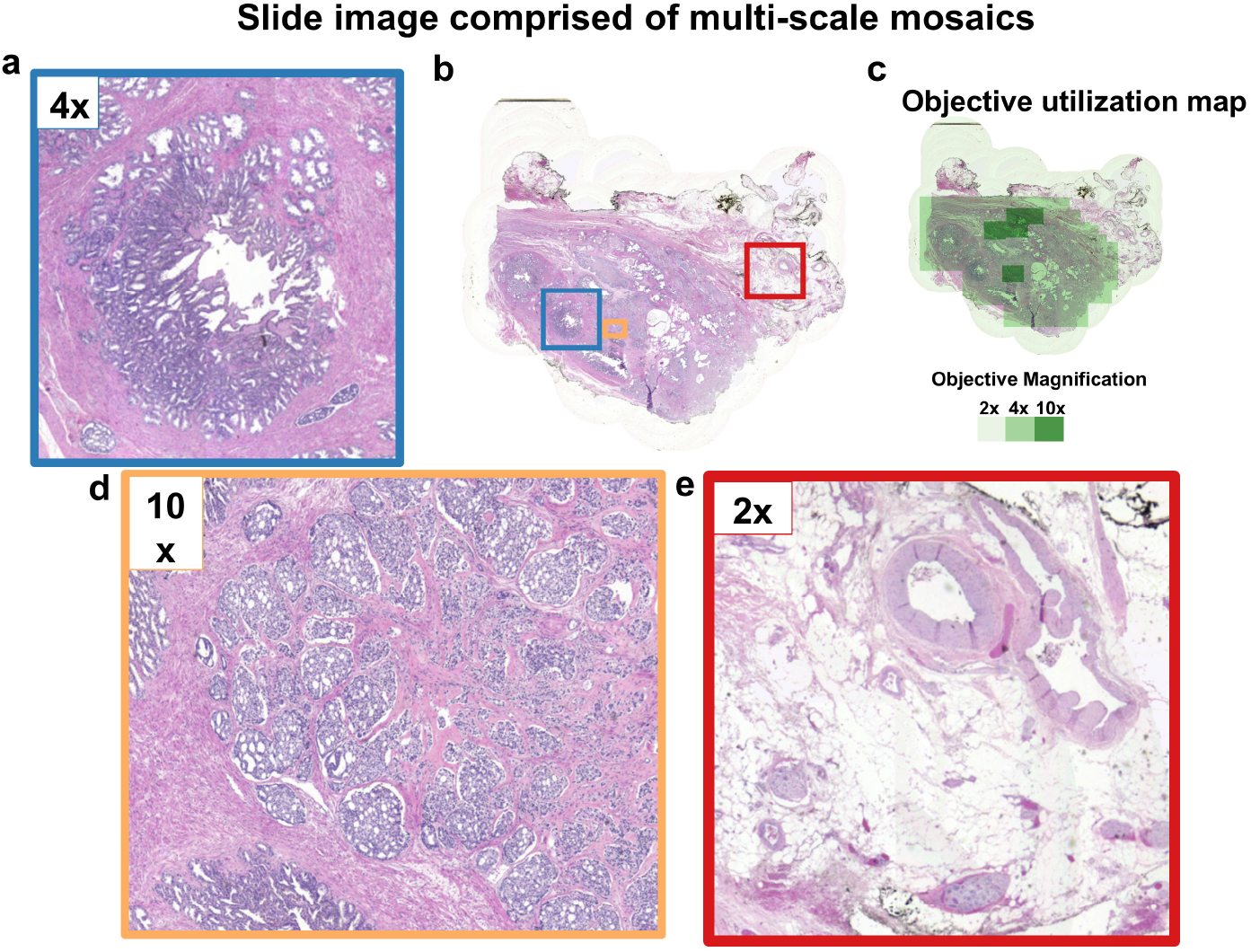
Multi-resolution slide image (MRSI) produced from glass slide review. This figure highlight areas of the final composite slide image **(b)** produced when the pathologist used different objectives **(c)**, including **a**, 4x, **d**, 10x, **e**, 20x objectives.

### 2.2 Data captured with PathCAM tell the story of the glass slide diagnosis

A unique and novel aspect of our approach is that, in addition to producing multi-resolution digital slide images, the registration processes used during video-mosaicking give us the spatial relationships between frames and, therefore, the location and scale of the ocular FOV over time. These previously uncaptured data can be used to visualize the spatiotemporal search processes that contributed to the pathologist’s diagnosis in clinical practice. Figure 5 shows an example of the datasets that were generated from a single recording session of a pathologist conducting a clinical manual glass slide review. Additionally, Supplementary Video 3 shows the original video recording alongside the pathologist’s path of analysis across one slide. In addition to the static MRSI (Figure 5a) and the associated magnification or resolution maps (Figure 5b), which represent which areas of the slide were viewed at what magnification, additional spatiotemporal data are collected that allow unique insights into the process of the review. For instance, timestamps and registration coordinates for each frame allow us to see not only where the pathologist looked and at what magnification, but also for how long. The latter data can be represented in a dwell time heatmap over the specimen area as in Figure 5c. Additionally, the overall path and sequence of the analysis can be reconstructed, as in Figure 5d, where the path of analysis is overlaid onto the PathCAM MRSI and is color-coded according to the temporal sequence and objective lens used (shown in the legend of Figure 5d).

**Fig 5.**
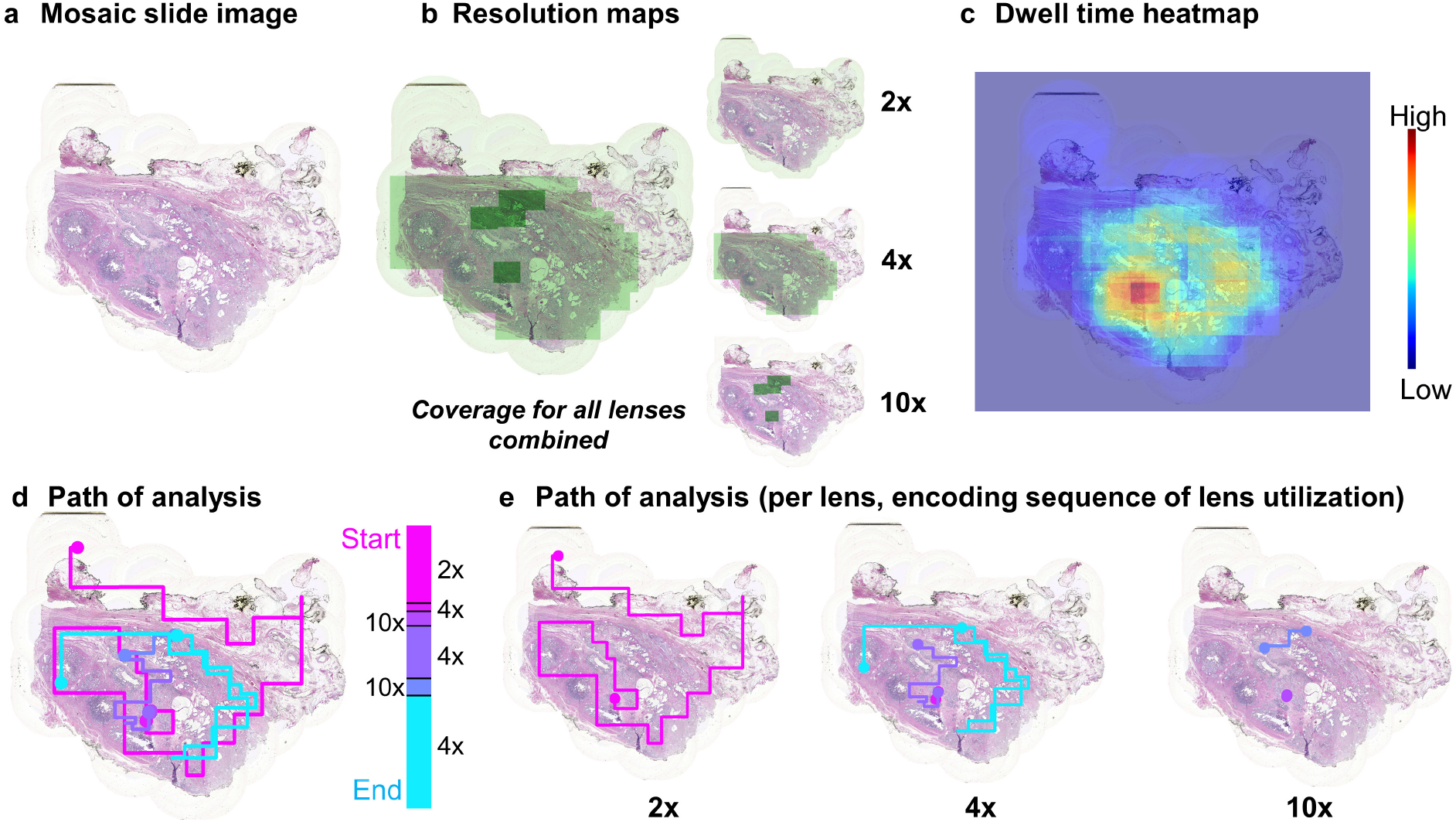
Novel data sets based on visual analytics of the pathologist at the microscope. **a**, Image data collected during the glass slide review are assembled into a single slide image. In addition, spatiotemporal data is collected, which comprise maps of **b**, utilized resolution, **c**, dwell time per pixel, and the path of analysis, both **d**, globally, and **e**, per lens used.

Additionally, the search path data can be isolated for each objective lens used, enabling one to observe the search strategy of the user as a function of magnification level (Figure 5e). These rich spatiotemporal data (Figure 5), combined with the image data present in the MRSI and in each captured frame itself, tell the story of which areas were most salient to the diagnosis, and the exact sequence of events contributing to the diagnosis. Such data, acquired from the gold standard of pathology diagnostic workflows, will be invaluable for a) the study of factors contributing to the accuracy and variability of pathologic diagnoses and b) the generation of copious amounts of image and user-interaction data useful for the development of a multitude of artificial intelligence approaches.

A critical aspect of our approach is that the user looks at the slide specimen through the microscope oculars at all times. Thus, the data captured with PathCAM are a true reflection of the pathologist’s natural behavior. Schaumberg et al. developed a method of registering microscope video frames to digital WSIs to determine the total dwell time across WSIs.^20^ However, magnification utilization patterns were not analyzed or visualized, and WSIs were still required to reconstruct the dwell-time heatmaps. Likewise, the commercially available microscope slide scanning system ‘Panoptiq’ by ViewsIQ allows multiple magnification levels to be recorded but requires the pathologist or operator to monitor the computer screen instead of conducting their normal clinical workflow.^18^ There have also been no studies that analyze the utilized magnification levels for this system. Other methods of scanning slides at the microscope using video-mosaicking only considered a single magnification level, which does not reflect the pathologist’s typical multi-objective workflow.^10–17, 19^ There are numerous studies that characterize user interaction with digital slides captured using WSI systems via multi-resolution viewers^21, 22^ or eye-tracking.^23–26^ However, to our knowledge, there are no technologies that record or visualize magnification levels, paths of analysis, and dwell time of pathologists reviewing glass slides at the microscope using unaltered diagnostic routines.

### 2.3 Pathologist review of digital multi-resolution slides constructed from glass slide analyses

One practical use of the digital slide images generated from the PathCAM approach in the clinical workflow is for the pathologist to double-check their work or send the digital images to a pathologist colleague for a quick second opinion review. To generate the multi-resolution slide images and test our approach, we asked an expert pathologist with genitourinary training to review a selection of 7 radical prostatectomy specimens from a single case. Radical prostatectomy was chosen because it represents a use case where specimen sizes and slide numbers per case are large, and a number of varied tasks are required during the review that have different demands on specimen search and magnification/resolution required (i.e., determination of overall tumor grade, overall tumor volume, determination of positive margin status, and detection of extraprostatic extension or seminal vesicle invasion (SVI)). Therefore, radical prostatectomy is an ideal test case for evaluating the capabilities of PathCAM. During the glass slide reviews, the pathologist determined the slides were mostly 3+4 Gleason grade. Intraductal carcinoma and extraprostatic extension were not identified. The pathologist determined that seminal vesicle invasion and perineural invasion were present. The primary tumor was described as PT3b, which indicates the tumor has invaded the seminal vesicle. During the glass slide reviews, the pathologist did not change their behavior to view or monitor the digital image acquisition. Therefore their diagnostic process was unaffected by the operation of the PathCAM system.

After processing the microscope recordings to generate the multi-resolution slide mosaics, we asked the same pathologist to review the produced digital images using the HistomicsUI viewer platform. For radical prostatectomy specimens, the pathologist would typically intend to cover the whole specimen area using the microscope, but have the limitation that they can only see one relatively small region of the slide at any given time, as their FOV is restricted by the FOV of the particular objective lens used. The maximum ocular FOV is 50 mm^2^ (using the 2x objective lens), whereas the size of the radical prostatectomy specimens used in this study was on the order of 135 mm^2^. Because the entirety of the specimen area in the digital PathCAM MRSIs generated during the slide review can be viewed within a single viewport on the user’s computer screen in contrast to the microscope, the PathCAM MRSIs provide additional spatial context to help the pathologist confirm whether they covered the entire specimen area, or whether any areas would require additional scrutiny. We visualized the completeness of a pathologist’s slide analyses in terms of the area of the specimen covered. During a recording session of 7 slides (Figure 6a), the pathologist covered 100% of the specimen area for 6 out of 7 specimens (Figure 7). For one specimen, 97% of the specimen was imaged during the original glass slide diagnosis. Less than 3% of this particular tissue specimen was missing in the final image because the pathologist did not pass over the center of the specimen. We confirmed that the gap in coverage was not due to any post-processing methods and was due to the fact that the pathologist did not pass over this small area during the diagnostic review. During a subsequent digital viewing session of the slide image captured during the glass slide review, the pathologist observed the missing area, and re-reviewed the glass slide to inspect this area (Figure 6d, 6e). This session was also recorded and was added to the original data stream to update the slide image, where it was confirmed that this subsequent recording session, combined with the first, resulted in a complete image of the entire specimen area (Figure 6c). Despite not covering a small area of the slide during the initial diagnostic review, the clinical diagnosis and impression did not change after reviewing this area of the glass slide during the second recording session. The only potential impact of not viewing the entire slide, in this case, would have been if the missing area had contained sufficient Gleason 4 to upgrade the slide from Gleason 3+4 to Gleason 4+3, or if there had been a small area of Gleason pattern 5 present, neither of which was the case. However, this result demonstrates an important potential feature of PathCAM, in that its real-time use could enable pathologists to check the completeness of their slide search and catch any potential diagnostic errors due to incomplete slide coverage or identification of important areas that received only cursory attention before they occur. As described in the previous section, besides generating an image that can be used for secondary review, our method captures the pathologist’s spatiotemporal interaction with the specimen, which can also be reviewed retrospectively as a useful platform for studying diagnostic variability within or across expert pathology reviewers.

**Fig 6.**
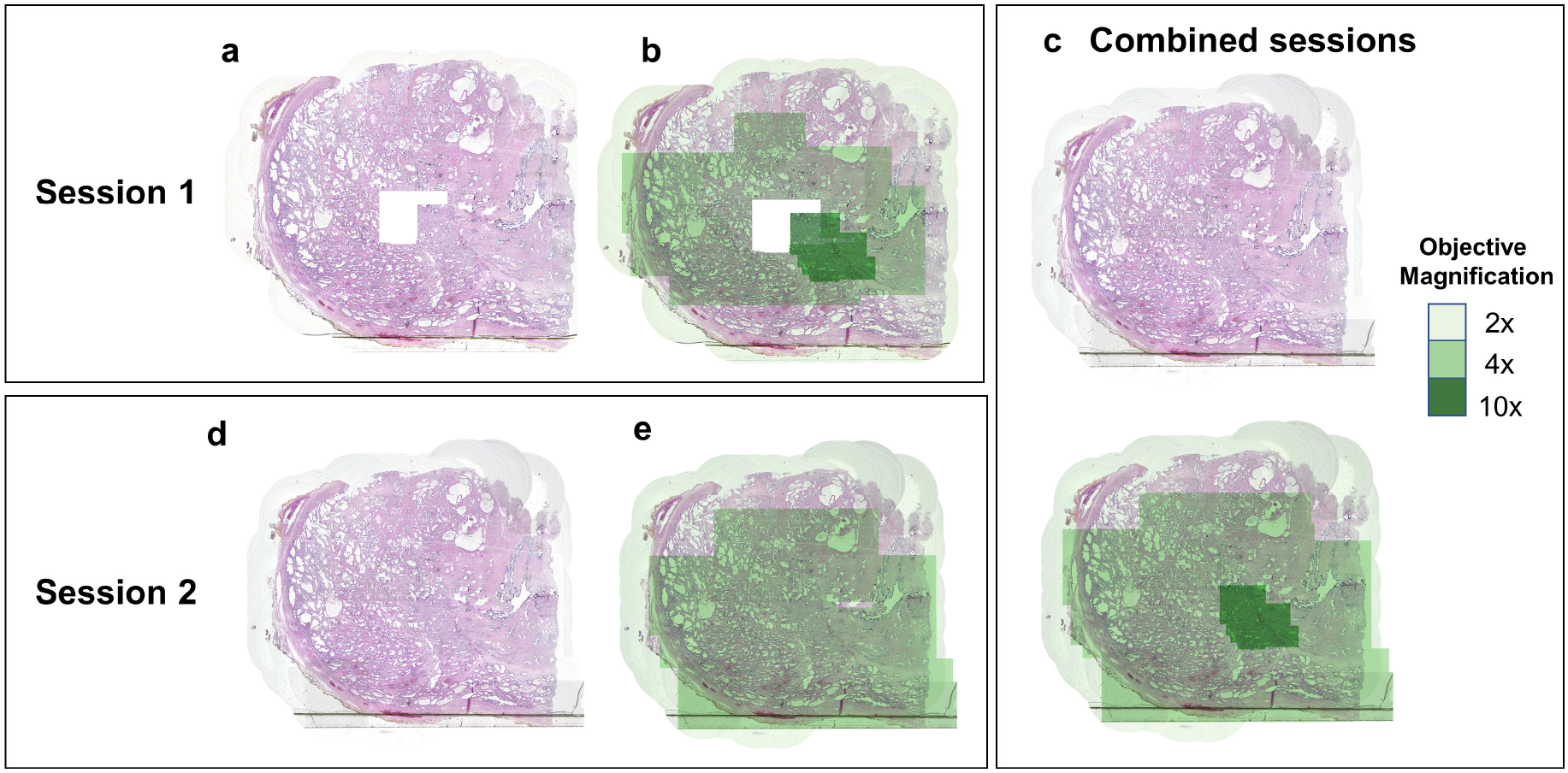
Combined multi-resolution slide image from multiple viewing sessions. **a**,**b**, This slide was not fully imaged during the first microscope recording session because the pathologist did not pass over the center of the slide. **d**,**e**, During the second microscope session, after reviewing the PathCAM MRSI, the pathologist subsequently covered the entire slide with the 2x and 4x objectives. **c**, Separate slide analysis sessions were merged to provide the updated digital record.

**Fig 7.**
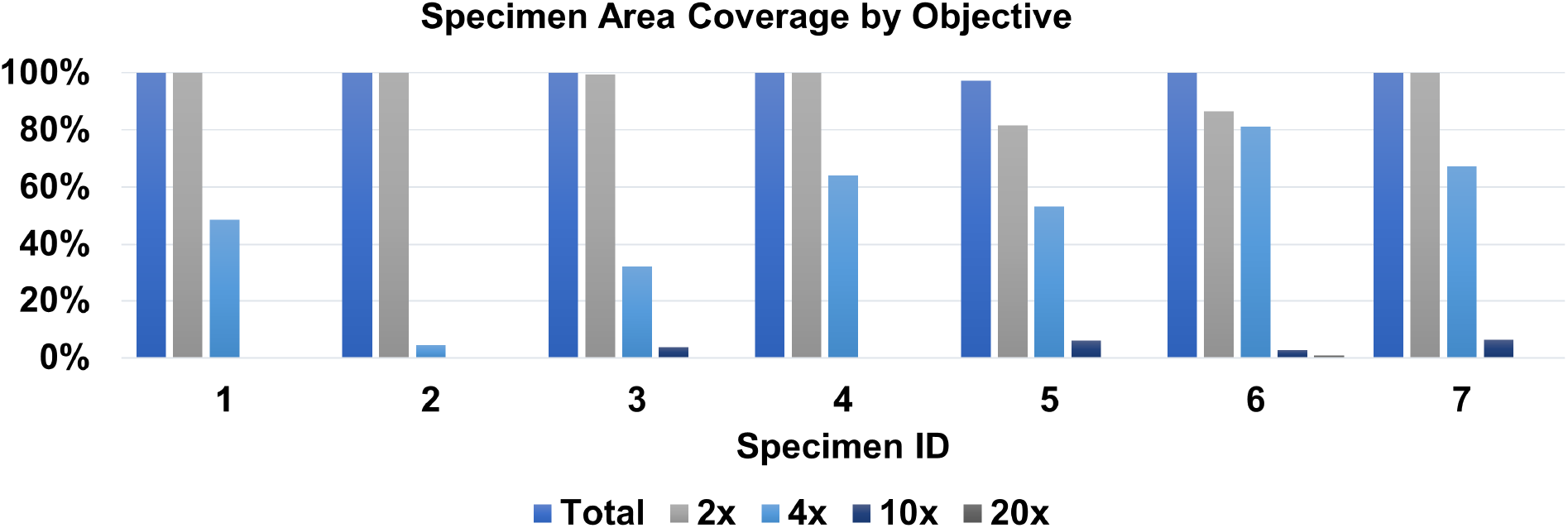
The percentage of specimen area imaged under the objectives.

### 2.4 Area and resolution coverage across the specimen areas

Besides creating a digital slide image during the microscope workflow, the mosaicking process also reveals important information about how the pathologist analyzed the slides. By capturing the exact movements of the pathologist at the microscope, we can visualize how the pathologist utilized objective lenses across the specimen area. We found that for this subset of radical prostatectomy slides, the pathologist mainly used low magnification objectives (2x and 4x) to review the majority of the specimen areas, and reviewed smaller areas with higher power objectives (Figure 7). The pathologist only used the 20x objective on one slide, which displayed seminal vesicle invasion. This is an interesting result because radical prostatectomy specimens are typically scanned with slide scanners at 20x or 40x magnification for clinical diagnosis on digital images,^27^ which appears to be unnecessarily high for the majority of the specimen area. Analyzing the pathologist’s magnification utilization may highlight which slides are most relevant to the diagnosis, or may even be useful to provide automated labels for regions of importance.

### 2.5 Application of digital annotation tools on PathCAM MRSIs captured during glass slide review

Although the primary diagnosis in this approach is conducted on the glass slide using the microscope as is standard of care, PathCAM provides digitized slide images to pathologists after the standard glass slide review, which gives them the added capability of utilizing digital tools, such as digital annotation. For radical prostatectomy specimens, for instance, tumor volume calculation could benefit from immediate access to the digital slide images. The typical approach for (analog) slide annotation using the microscope is for the pathologist to draw on the slide using an ink marker, for example, drawing dots around the tumor volume, indicating the location of positive surgical margins, or providing other text labels, among others. These physical ink annotations may be subsequently used for calculation purposes. For instance, after removing the slide from the microscope, the pathologist may estimate the tumor volume by visually sizing up the outlined areas. While this is the standard practice, it is clear that digital annotation could help the pathologist computationally quantify tumor volume, outline areas of the tumor, and provide digital labels on areas of interest. Figure 8d shows an example of the pathologist’s digital tumor volume annotation (black) on the final slide image. Additionally, the PathCAM MRSIs could be used for digital analysis such as computational nuclear or glandular segmentation (Supplementary Figure 7) or other computational assists as desired. The full range of utility of PathCAM MRSIs for computational analysis will be investigated in future work.

**Fig 8.**
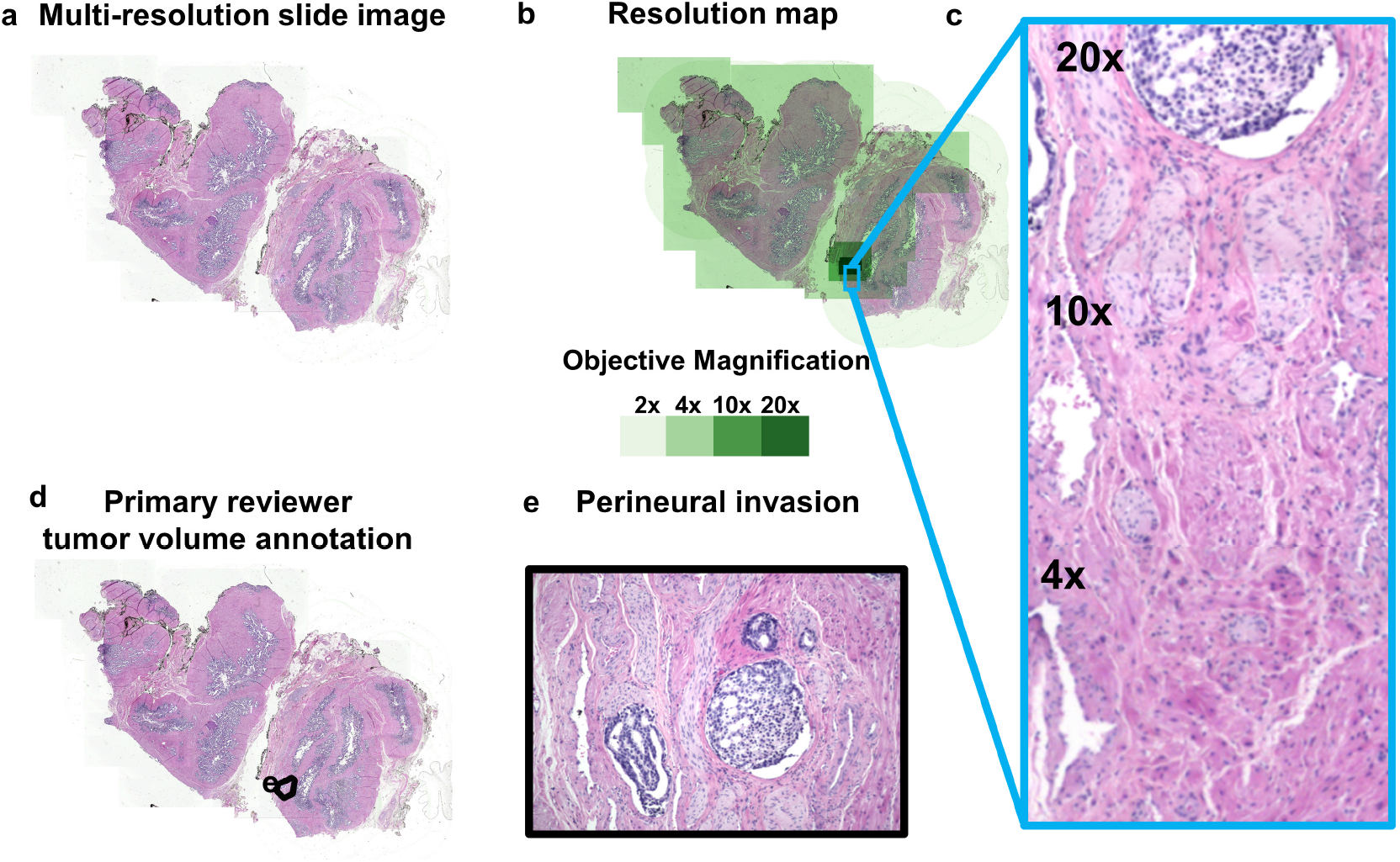
Demonstration of tumor annotation on MRSI during digital review, integrating digital annotation into the glass slide review. **a**, The multi-resolution slide image produced from the microscope slide review. **b, c**, The ROI was imaged with 20x magnification by the pathologist. **d**, The pathologist annotated (black outlines) the tumor area using HistomicsUI. **e**, This inset shows the location of perineural invasion.

## 3 Conclusions and Future work

The proposed solution outlined in this paper is based on a simple insight that the video feeds from cameras installed on clinical microscopes can be used to build complete digital records of clinical slide reviews. Unlike previous methods, we did not involve the pathologist user directly in image generation, but rather “ambiently” generated digital images of the specimen that were directly reflective of their natural processes and then investigated the potential utility of those images to be used in a feedback loop to improve the glass slide diagnosis. The data produced by PathCAM are reflective of the information leveraged by the pathologist user to render a diagnosis on the glass slide - as such, PathCAM MRSIs may not be a one-to-one replacement for *de novo* primary digital diagnosis using standard WSI. However, the PathCAM approach provides a number of attractive features, based on its human-in-the-loop nature, to bridge the gap between glass slide microscope diagnosis and digital pathology capabilities, that will make it a valuable complement to WSI. This study confirms the feasibility of this onboard-camera approach as a novel, low-cost, hybrid analog-digital pathology platform that could enable disruptive innovations in both clinical pathology and digital pathology research.

This work demonstrates the potential of the PathCAM system to connect digital pathology capabilities to the current analog clinical pathology workflow and to provide novel and rich human-machine interaction datasets that could be tremendous resources for future research and clinical decision support. The ability to support both real-time computational assistance and fast digital collaboration to improve diagnostic accuracy in a larger dataset will be studied in future work. We will also study whether inter- and intra-observer variability could be reduced by providing the pathologist with the PathCAM MRSIs in real-time to double-check their diagnosis and solicit collaborative opinions.

Although the frame registration process can potentially be done while streaming, the frame sampling process is currently not done in real-time. Frame sampling is important to remove redundant frames and reduce overall data size while ensuring that the highest quality frames that ensure complete coverage are retained for accurate mosaic and spatiotemporal attention map reconstruction. To expedite data processing for real-time mosaic generation, we intend to explore more efficient methods of sampling the video stream to reduce data overhead. While the fixed subsampling ratio used here provides gap-free mosaics, there are still areas that are oversampled. Although video-recording the pathologist at a fixed frame rate of 26 FPS led to oversampling the specimen area, we were able to reduce the number of frames used in the final mosaics by 93% *±* 1.5% using fixed subsampling. However, we were conservative in reducing the number of out-of-focus frames because these frames may cover areas of the specimen which are not covered by other frames. In other words, we prefer to include slightly out-of-focus frames rather than display a gap in the mosaic. We also observed that while most frames were in focus (note that the user controlled the focus), the focus was more likely to be slightly off when using the higher power objectives due to the smaller depth of field of the objective lenses and the eye’s ability to accept a wider range of focal conditions than the camera. In the future, we could artificially enhance the focus of these frames,^28, 29^ or use a more sophisticated technique such as graph cuts^30–32^ to reduce the overlap of out-of-focus frames over in-focus frames.

Finally, we will work to disseminate our inexpensive approach to enable image data to be captured widely from a more equitable variety of sources that do not have access to costly scanners. We will also evaluate the performance of machine learning models applied to the MRSIs to enable computer-aided diagnosis and data collection where traditional digital pathology is unavailable. One study found that most clinical machine learning algorithms were trained using data from patient cohorts from California, Massachusetts, and New York, which raises serious questions on model generalizability across disparate patient populations from diverse geographical locations.^33^ Application of models trained on specific patient populations may have lower performance when applied to new patient populations, or edge cases that they were not trained on.^34^ Our system would put digital imaging directly into the microscope workflow used on the vast majority of slides today, and for the foreseeable future. This would ultimately enable the collection of previously uncaptured data from diverse sources, thereby supporting the development of more generalizable AI-based computational decision support systems and enabling those systems to be used in realtime during the standard clinical glass slide review. With the PathCAM approach, a greater fraction of pathologists and patients could have access to these emerging digital pathology technologies, which are expected to make a significant impact on improved diagnosis and prognosis.

## 4 Methods

### 4.1 System design

To capture the pathologist’s microscope analyses, we attached a large format CMOS camera (Sony IMX342 / FLIR Oryx Color 31MP) on a Nikon Eclipse Si upright microscope equipped with 2x (Nikon CFI60 Plan Achromat Ultra Wide 2x Objective Lens, NA 0.06, WD 7.5 mm, F.N. 25 mm), 4x (Nikon CFI60 E Plan Achromat 4x Objective Lens, NA 0.1, WD 30mm, F.N. 22 mm), 10x (Nikon CFI60 E Plan Achromat 10x Objective Lens, NA 0.25, WD 7mm, F.N. 22mm), 20x (Nikon CFI60 Plan Achromat 20x Objective Lens, NA 0.4, WD 1.2mm, F.N. 22mm), and 40x (Nikon CFI60 E Plan Achromat 40x Objective Lens, NA 0.65, WD 0.65mm, F.N. 22mm) objectives. The microscope’s light source was split to provide 50% to the eyepieces (Nikon CFI 10x Eyepieces, F.N. 22mm) and 50% to the camera’s sensor using a Nikon C-TT trinocular tube.

#### 4.1.1 Camera Spatiotemporal Sampling

We chose a camera that both properly sampled the optical resolution of the microscope at all magnifications, satisfying the Nyquist-Shannon criterion,^35^ and that also provided sufficient temporal sampling to accommodate unconstrained slide movement during analysis. The Nyquist-Shannon criterion requires a camera’s pixel size to be *at least* half of the smallest resolvable optical dimension to preserve resolution at the sample. This means,

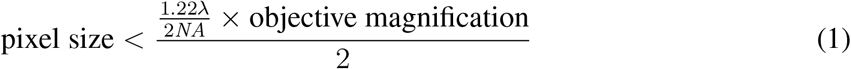

where NA is the numerical aperture of the objective and *λ* is the wavelength of light. For each objective and numerical aperture (NA), the largest pixel size allowable to maintain the optical resolution is shown in Table 1. With our current camera sensor (Sony IMX342 Color, 6464 × 4852 pixels), the pixel size is 3.45 microns. We verified that at each of the provided objective magnification levels (2x to 40x), the pixel size of the camera is at least a factor of 2 smaller than the maximum pixel size, satisfying the Nyquist-Shannon sampling requirement (shown in the last column of Table 1).

**Table 1.** Camera resolution requirements to preserve optical resolution. For each objective magnification level, we calculated the theoretical resolution limit and projected size based on the Nyquist criterion. The maximum pixel size is the projected size divided by 2. Our camera sensor pixel was less than the maximum pixel size across all objective magnification levels.

| Magnification | NA | Resolution limit (um) | Projected Size (um) | Maximum pixel size (μm) | Camera sensor pixel size (μm) | Pixel over-sampling factor |
| --- | --- | --- | --- | --- | --- | --- |
| 2 | 0.06 | 5.59 | 11.2 | 5.59 | 1.73 | 3.24x |
| 4 | 0.1 | 3.36 | 13.4 | 6.71 | 0.86 | 7.78x |
| 10 | 0.25 | 1.34 | 13.4 | 6.71 | 0.35 | 19.45x |
| 20 | 0.4 | 0.84 | 16.8 | 8.39 | 0.17 | 48.62x |
| 40 | 0.65 | 0.52 | 20.6 | 10.32 | 0.09 | 119.69x |

In order to passively capture slide images based on the natural viewing pattern of the pathologist, it was important for the camera’s FOV to cover what the pathologist could see through the microscope ocular, while also maintaining sufficient sampling resolution. We chose not to use a de-magnifying camera adapter, which would increase the field of view at the expense of resolution. The size of the ocular field of view is the field number (F.N.) of the oculars divided by the objective magnification. We calculated that the diameter of the field of view of view through the oculars was 5.5 mm, 2.2 mm, 1.1 mm, and 0.55 mm at 4x, 10x, 20x, and 40x magnification, respectively. The ocular FOV coverage of the camera sensor using the 4x, 10x, and 20x objectives was therefore 86%. Because the F.N. of our 2x objective (25 mm) was larger than the F.N. of the oculars (22 mm) and microscope tube diameter (22 mm), the captured image at 2x magnification was vignetted on the camera sensor and the ocular field of view was reduced from 100%. Due to vignetting, the darkened edges of the images with the 2x objective suffered from low signal-to-noise ratio (SNR) after flat field correction. As a result, we cropped these frames, ending up with a FOV coverage of 74% of the full ocular FOV at 2x magnification.

#### 4.1.2 Camera frame rate

We determined the required camera frame rate according to the maximum stage expected translation speed in combination with the frame size. The camera had to acquire images quickly enough such that there were no gaps in the final stitched mosaic due to excessive inter-frame movement of the slide. The frame rate had to be high enough to ensure a minimum level of overlap between adjacent frames at maximum stage speed, which we conservatively set as 20% for the SIFT algorithm. We measured maximum expected stage speed (7 mm/s) by recording a pathologist reviewing radical prostatectomy specimens with a high frame rate (200 FPS) camera (FLIR Blackfly S, CMOS, 0.4 MP) (Supplementary Figure 1). We found the displacement between successive frames in the video stream using SIFT/RANSAC, described in detail below, then calculated stage translation speed based on the frame’s millisecond-resolution timestamp. The minimum required frame rate is then a function of stage speed. The maximum stage translation displacement which maintains at least 20% overlap is given by the equation:

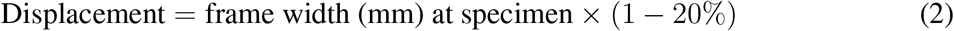

Therefore, the minimum frame rate to ensure the specified overlap is:

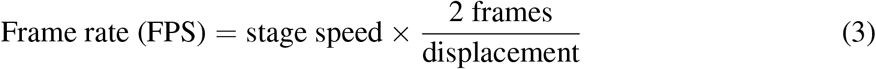

Using these equations, for our chosen camera sensor (Sony IMX342, APS-C format), we found that the minimum frame rate to ensure a gap-free video stream using all objective lenses was 7 FPS, well below the maximum full-frame speed of the FLIR Oryx 31MP camera of 26 FPS. To allow for redundancy, we set the frame rate to the camera’s maximum, 26 frames-per-second (FPS) using image acquisition software (Spinview, Teledyne FLIR) and a Dell Precision Tower 5810 with 64 GB RAM and Nvidia NVS 300 GPU. Through the acquisition software, we set the exposure mode to automatic and set the upper bound of the exposure time to 1000 *µ*s. We chose a camera with a global shutter as opposed to a rolling shutter to ensure each pixel in the sensor was exposed simultaneously, meaning the captured frames would not appear distorted when the specimen moved. Video frames were captured in Bayer RG8 Raw and converted to TIFF during post-processing. The resolution of each frame was 4852 × 6464 pixels.

### 4.2 Coarse registration

After the recording sessions, we converted the captured RAW frames to TIFF. All processing steps are shown in the diagram Figure 9. The video frames were matched by relating one image’s features to a second image’s corresponding features, using the feature detection algorithm “Scale Invariant Feature Transform” (SIFT). This algorithm extracts features from images at different scales, making it robust to differences in magnification and frame focus variations.^36^ We used OpenCV’s^37^ implementation of SIFT to detect features after rescaling the images to a width of 800 pixels. After features were detected, the features from one image were paired with the features of the next image using brute force k-nearest neighbor matching (k=2). False positives matches were pruned with Lowe’s ratio test,^36^ keeping matches only if the ratio of the Euclidean distances between a keypoint’s first and second matching keypoints was above 0.75.

**Fig 9.**
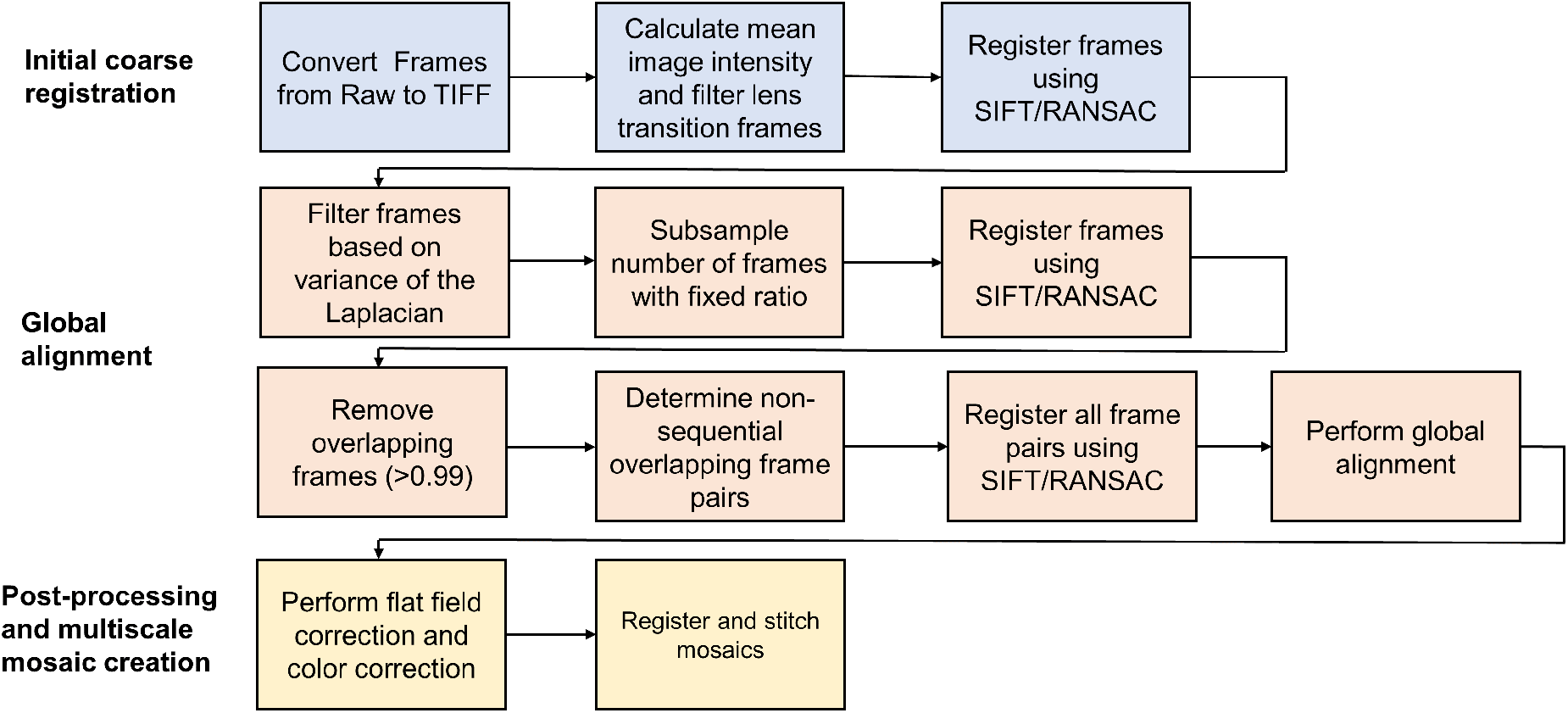
Overview of mosaicking process. To create the final mosaics, we performed a number of processing steps on the raw frames data to reduce visual misalignments and correct the color of the frames.

Because the glass slide on the microscope stage and camera sensor planes were roughly parallel, a 2D approximation was used, and no rotation was assumed. The Random Sample Consensus (RANSAC)^38^ method was used to estimate the best set of matches iteratively. A random selection of four corresponding points was selected during each iteration, and the transformation matrix was calculated. The number of inliers, or correspondences where the residual was less than two pixels, were counted. After repeating the random selection and transformation calculation process 75 times, the set of matches with the largest number of inliers was kept, and the final 2D transformation was calculated.^39^ After calculating the translation between the two images and the process was repeated for all frames.

### Global alignment and post-processing

For all frames, we calculated mean image intensity and the variance of the Laplacian, the latter as a measure of focus.^40^ We removed frames from the sequence with low measures of focus and image intensity, typically those with values greater or less than 2 standard deviations away from the mean and less than 100 matches, found using the RANSAC process as described above. These typically included frames acquired when the objective lens changed, efficiently removing those from the final video stream. Next, as a simple initial approach, we subsampled the number of frames by 10 to reduce the data stream. If gaps in the mosaic were detected, we reduced the subsampling factor to 5.

Because image mosaics are the cumulative result of the calculated translations between many successive image pairs, image registration errors accumulate as mosaics are built. Registration errors become particularly noticeable across looping paths, where non-sequential images overlap and any error in frame placement is obvious.^41^ To prevent this issue, other slide mosaicking methods utilized an automated stage with consistent acquisition patterns and recorded stage coordinates to aid in error minimization when aligning and stitching images.^10–12^ In contrast to techniques which require additional hardware, we optimally registered images by applying global alignment to the subsampled image sequence. Global alignment is a technique commonly used when constructing panoramas^42, 43^ and has been used in medical image mosaicking, such as in 3D in-vivo imaging.^44–46^ Global alignment adjusts the locations of images in a panorama so that global error across image positions is minimized. More specifically, global alignment is the linear least-squares minimization of the system of equations relating the locations of each overlapping image in the panorama (sequential and non-sequential) to the translations between all matched keypoints for the corresponding image pair. To increase computational speed, we simplified the global alignment problem by only including the final image translations *t*_*ij*_ = (*t*_*xij*_, *t*_*yij*_)^*T*^, found through RANSAC, in the model (Equation 4).

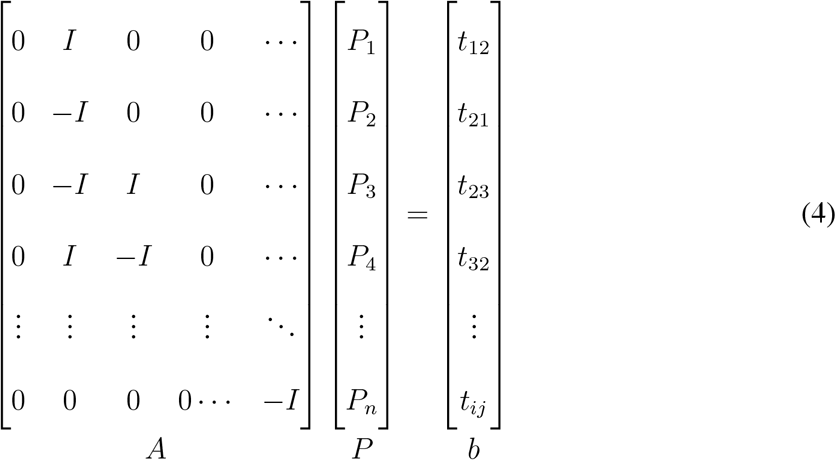

where **I** denotes the 2×2 identity matrix, **n** is the total number of frames, and **P** is the top-left co-ordinate of each frame, representing (*x*_*i*_, *y*_*i*_)^*T*^. The 2D translations, or Euclidean distances relating the images, are *t*_*ij*_. The first image was positioned at (0,0). The solution to minimizing ||*AP* −*b*||is:

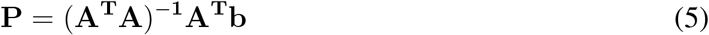

Solving Equation (5), returns the re-aligned coordinates of each image, **P**.

However, before applying global alignment, we further subsampled the sequence of frames based on percentage of area overlap of the frame to its neighbor. We computed the percentage of area overlap of each frame to the previous frame in the sequence. If frames overlapped by 99% or more, indicating the microscope stage was stationary, the second frame was removed. This eliminated excess frames covering the same area, such as when the stage remained stationary.

Next, to find overlapping frames through nearest neighbor search, a k-dimensional tree (k-d tree) was constructed where each node represents the top-left coordinate of each frame in the mosaic. SIFT and RANSAC were used again to find matches and translation between all overlapping image pairs, which we define as neighbors within 0.5 of the frame width of each node of the tree. To remove poor-quality matches, pairs were removed if the number of match inliers over total matches was less than 0.1. We then constructed the set of linear equations (Equation 4) of all paired image translations and solved for **P**. Supplementary Figure 8 shows an example of a mosaic before and after global alignment, which corrected a visible misalignment across a looping path.

The frames comprising the final mosaics were then flat-field corrected by dividing the frames by a blank calibration image taken with the corresponding objective. After finding the globally adjusted spatial relationship between all frames and correcting the frames, the frames were composited into mosaics. The white balance of the individual mosaics was corrected by finding the minimum and maximum intensity per color channel (in the range of 5 to 95%) and rescaling the pixel values to be within these limits for each color channel.

A typical PathCAM recording thus results in a series of globally-registered mosaics across multiple scales, reflecting the user’s choice of objective lens during the sequence of review. The adjoining ends of each of these mosaics were matched using the SIFT/RANSAC process to find the difference in scale and translation between mosaics.The final slide image was creating by transforming each mosaic into the space of the highest resolution. For instance, if the pathologist utilized the 4x and 2x objectives, all 2x mosaics were rescaled to 4x scale to compose the final multi-resolution slide image (MRSI) (Figure 3).

### 4.3 Computational analysis and tools

The final MRSIs were uploaded to the slide-viewing platform, HistomicsUI (https://github.com/DigitalSlideArchive/HistomicsUI). HistomicsUI allows the user to annotate images and provides a logging function to track zoom level, viewport size, and viewport location, among other functionalities. Supplementary Video 2 shows HistomicUI’s user interface and zooming function on one of our generated slides. The original image was comprised of multiple mosaics ranging from 2x to 20x magnification to create the mosaic image. As mentioned previously, we upsampled all mosaics to match the scale of the 20x mosaic.

We also visualized the dwell time of the pathologist’s digital and glass review by summing the cumulative time interval spent at each pixel and applying a color map to the images based on the maximum and minimum time intervals for each session (Supplementary Figure 6).^47^

### 4.4 Study design

We designed the pilot study to gather feedback on the feasibility of passively generating slide images from pathologist slide reviews and the potential utility of the resulting digital images and saliency data. We first conducted a recording session with an expert pathologist reviewing glass slides using the system described above. During the session, we recorded the pathologist reviewing one radical prostatectomy case consisting of a selection of 7 hematoxylin and eosin (H&E) stained slides. We chose radical prostatectomy slides to test the feasibility of our system due to the large size of the specimens and the need for various resolution levels across different diagnostic tasks. During each slide recording, the pathologist conducted several diagnostic tasks according to the standard radical prostatectomy College of American Pathologists (CAP) report, including identification of Gleason pattern, tumor margin status, presence of extraprostatic extension (EPE), invasion of the seminal vesicle, etc.^48^ Before the recording session, we calibrated the ocular focus to the camera focus by adjusting the diopter rings on the eyepieces. Once the recordings began, the pathologist exclusively looked through the microscope oculars without stopping to annotate slides, make notes, or view the recorded data. The pathologist moved the stage freely during the analysis. Post-analysis, the pathologist recorded diagnostic results according to the CAP report.

Approximately three months after the initial review of the radical prostatectomy case, the same pathologist reviewed the generated slide images through HistomicsUI. After referring to their previously recorded diagnostic information captured in the CAP report, the pathologist reviewed each digital slide for the same diagnostic features. Additionally, the pathologist was asked to annotate the tumor volume according to Gleason grade through HistomicsUI. The pathologist was allowed to refer back to the glass slides under the microscope, if desired, at any time while reviewing the diagnosis, which was also recorded (Figure 10).

**Fig 10.**
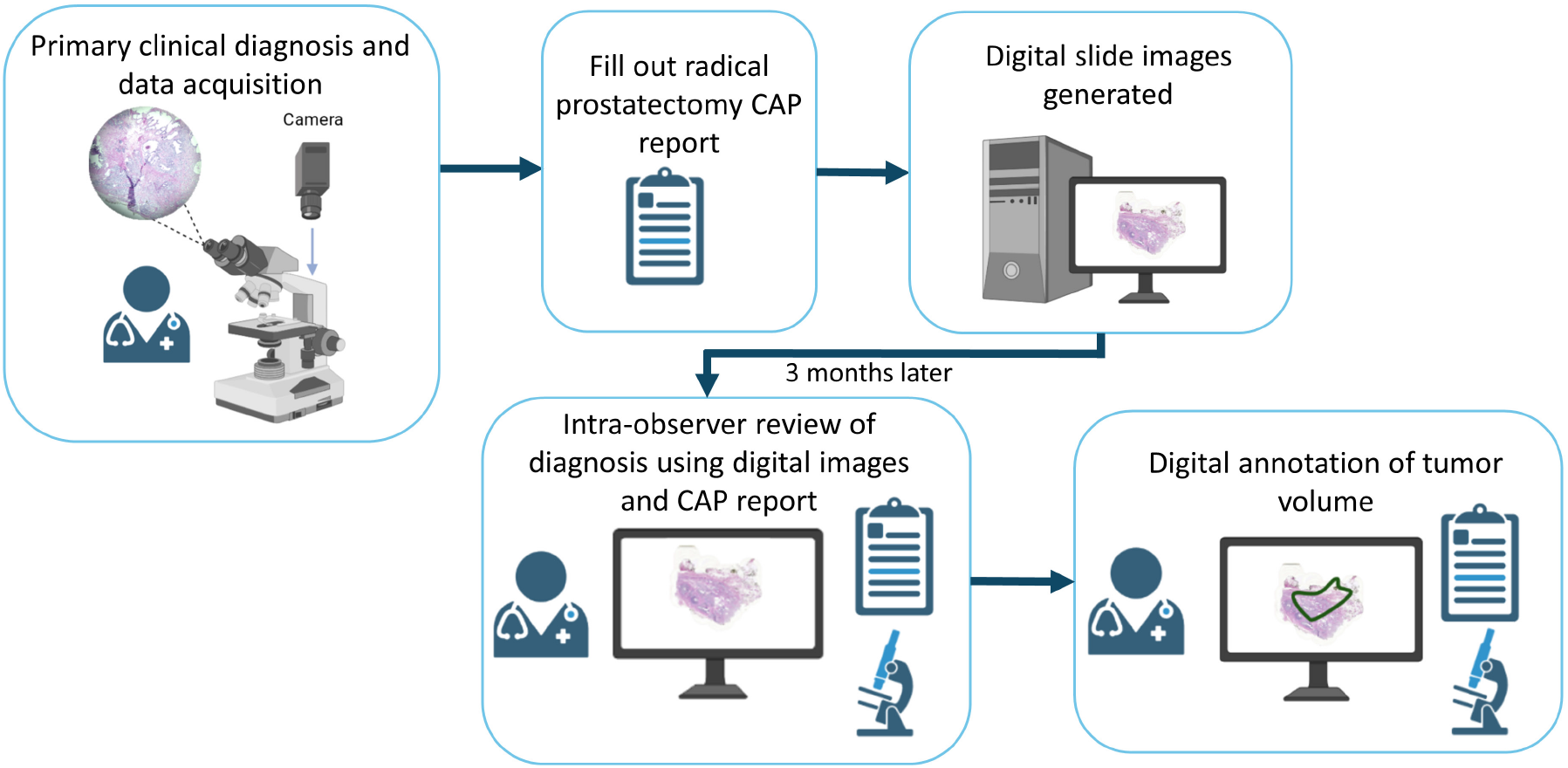
Study workflow. First the pathologist reviewed each slide at the microscope while the microscope camera was recording. After reviewing all slides, the pathologist recorded the diagnostic results in the CAP report. After generating the digital slide images, the pathologist reviewed the slides digitally. Additionally, when the pathologist was asked to digitally annotate the tumor volume, the pathologist had the opportunity to review the glass slides.

### Disclosures

JQB is a shareholder and employee of Instapath, Inc., which did not financially or materially support this work. No other authors have disclosures to report.

## Acknowledgments

Research was supported by Award Number 1664848 awarded by the National Science Foundation - Division of Mathematical Sciences, Award Number 2136744 awarded by the National Science Foundation - Division of Information & Intelligent Systems, Award Number 1144646 awarded by the National Science Foundation – Division of Graduate Education, and Award Number R01CA222831 awarded by the National Institutes of Health, National Cancer Institute. We would also like to acknowledge David Manthey and Roni Choudhury of Kitware for their support in implementing HistomicsUI.

## Supplementary Information

**Supplementary Figure 1.**
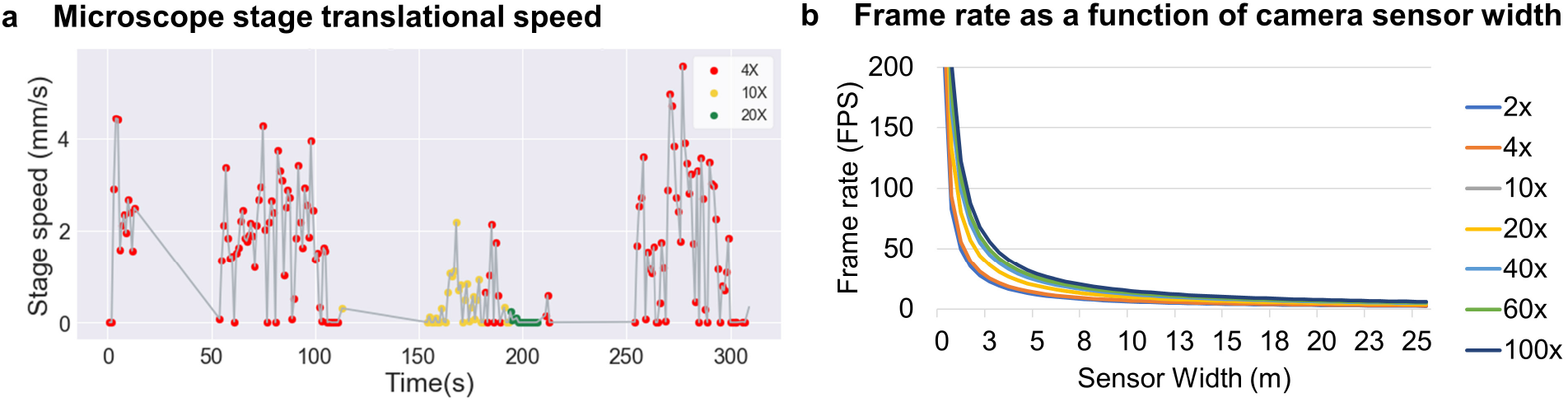
Maximum measured stage translation velocity and corresponding frame rate as a function of camera sensor width to maintain coverage at maximum stage velocity. **a**, Using a small-format high-frame rate camera (FLIR Blackfly S, 200 FPS) and video-mosaicking to calculate the translation between frames, we measured the speed of the stage as the pathologist reviewed specimens. The maximum stage translation speed was approximately 7 mm/s. **b**, Required frame rate as a function of camera sensor width for all objective lenses to ensure gap-free coverage at maximum stage velocity. The Sony IMX342 sensor selected for the PathCAM prototype has a width of 22.3 mm, and the minimum frame rate to ensure overlap using all objective lenses (2x to 100x) was approximately 7 FPS, well below the maximum 26 FPS available with this camera.

**Supplementary Figure 2.**
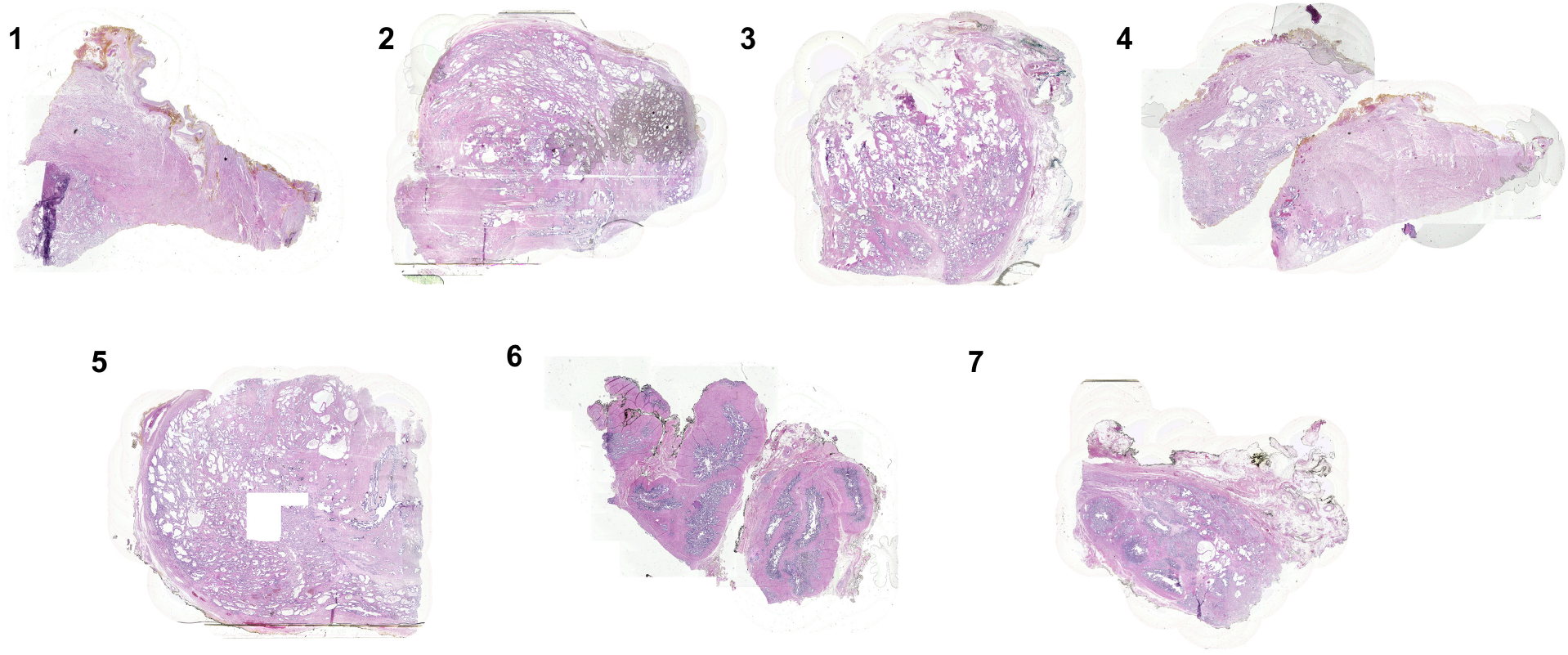
Multi-resolution slide images produced from microscope review. MRSIs generated from multiscale pathologist slide reviews without altering the diagnostic behavior of the pathologist.

**Supplementary Figure 3.**
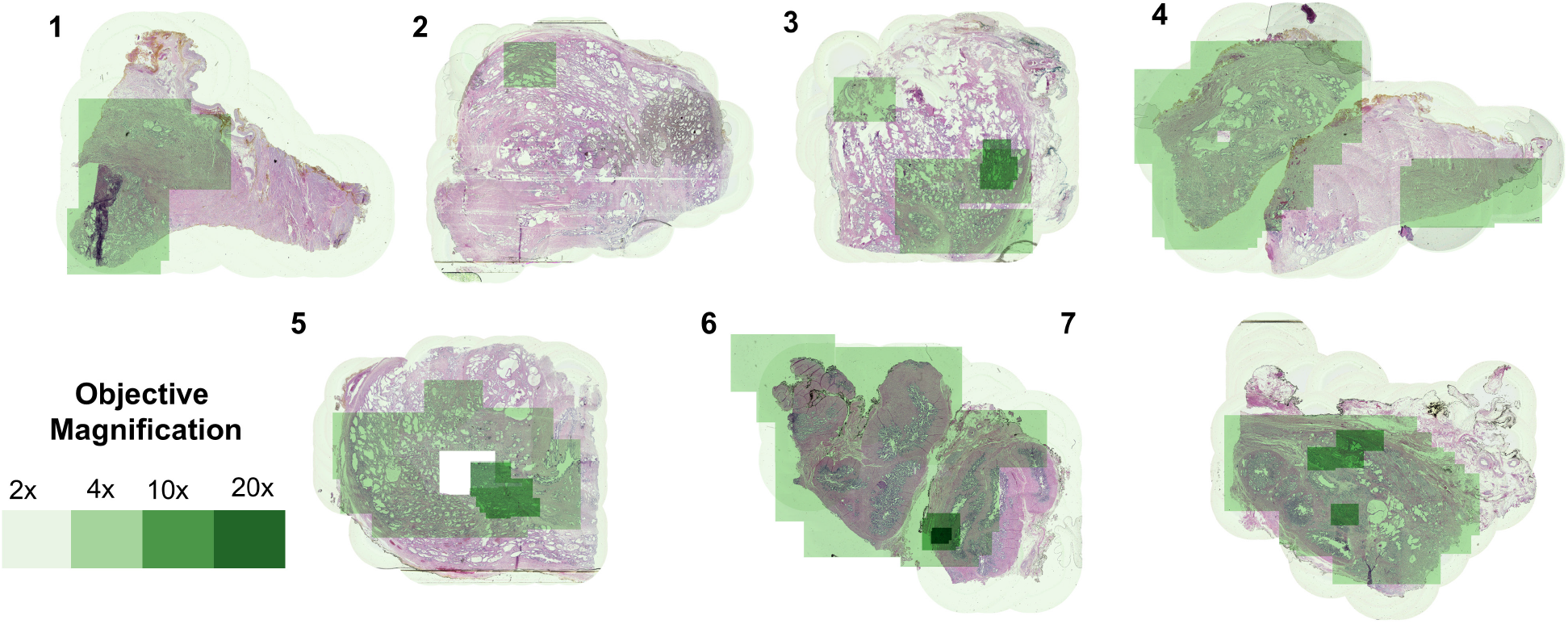
Resolution maps showing objective magnification levels used by the pathologist at the microscope. The green overlays indicate the objective magnification (4x to 20x) used by the pathologist across the specimen area. Darker green overlays indicate higher objective magnification.

**Supplementary Figure 4.**
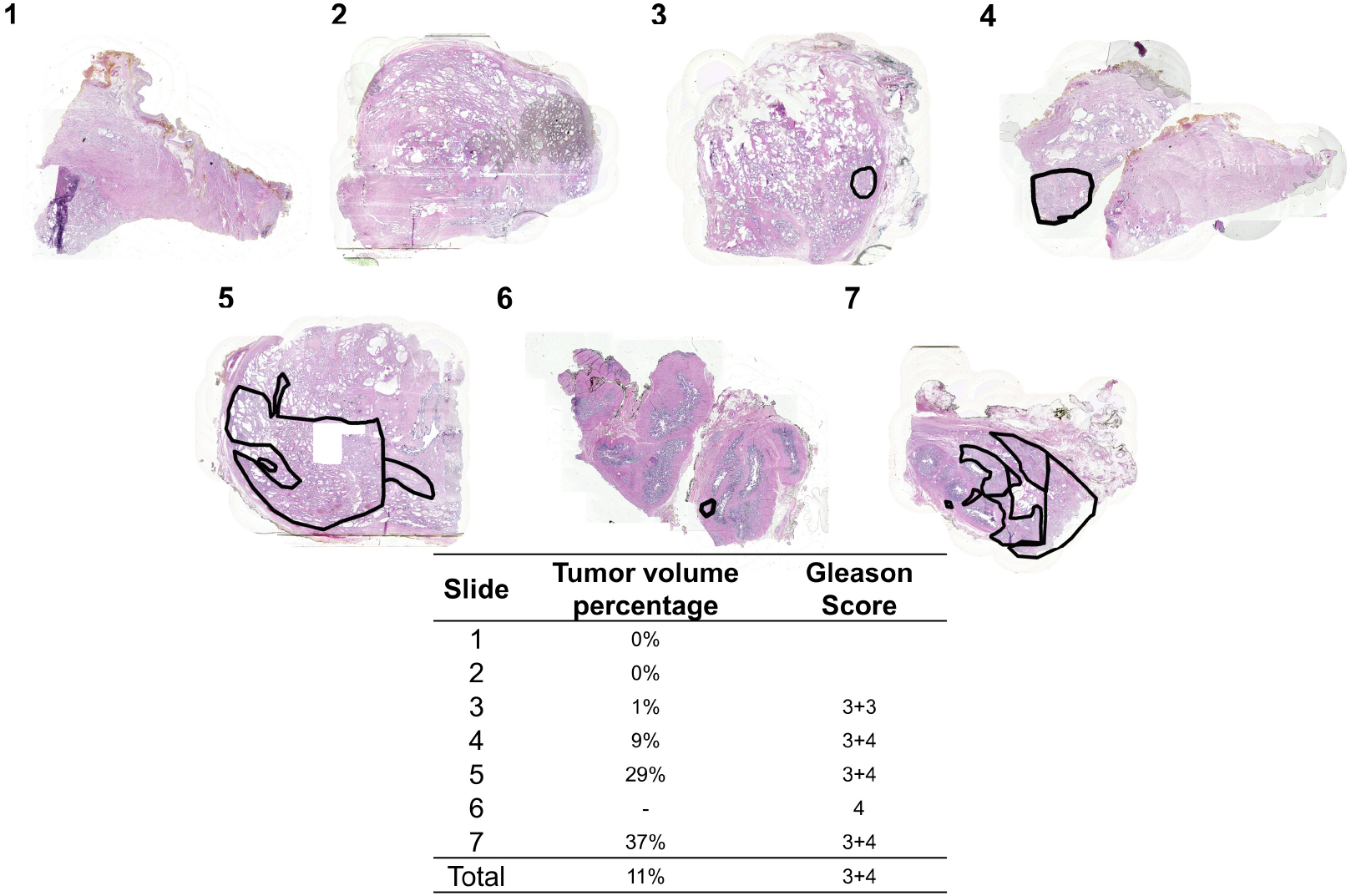
Digital tumor volume annotations of multi-resolution slide images. During the MRSI review session, the pathologist annotated the tumor volume (black outlines) in HistomicsUI. Slides 1 and 2 do not contain any tumor. Slide 6 was the seminal vesicle specimen and was not included in the tumor volume calculation.

**Supplementary Figure 5.**
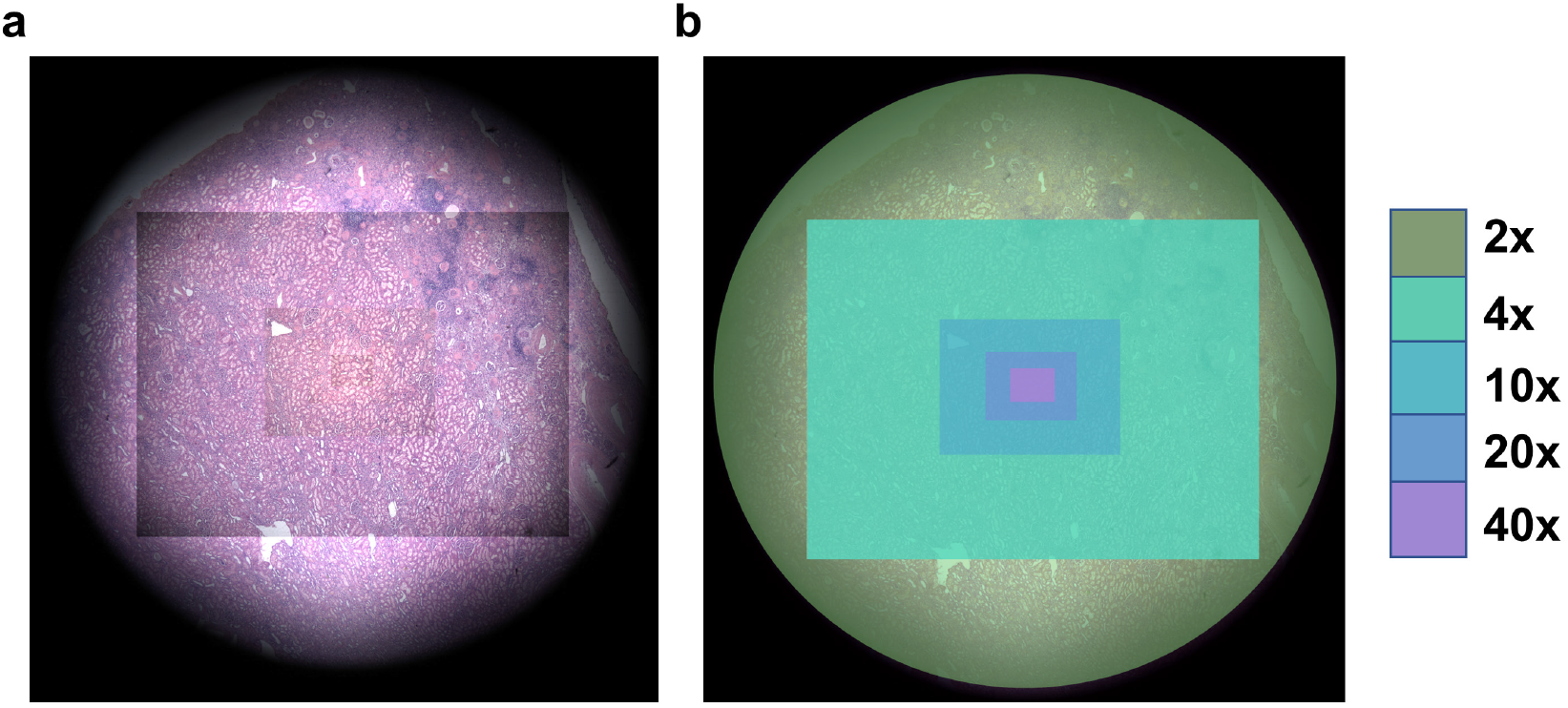
Camera FOV (APS-C camera FOV). **a**, Registered images from each lens at a single location. **b**, Camera FOV per lens

**Supplementary Figure 6.**
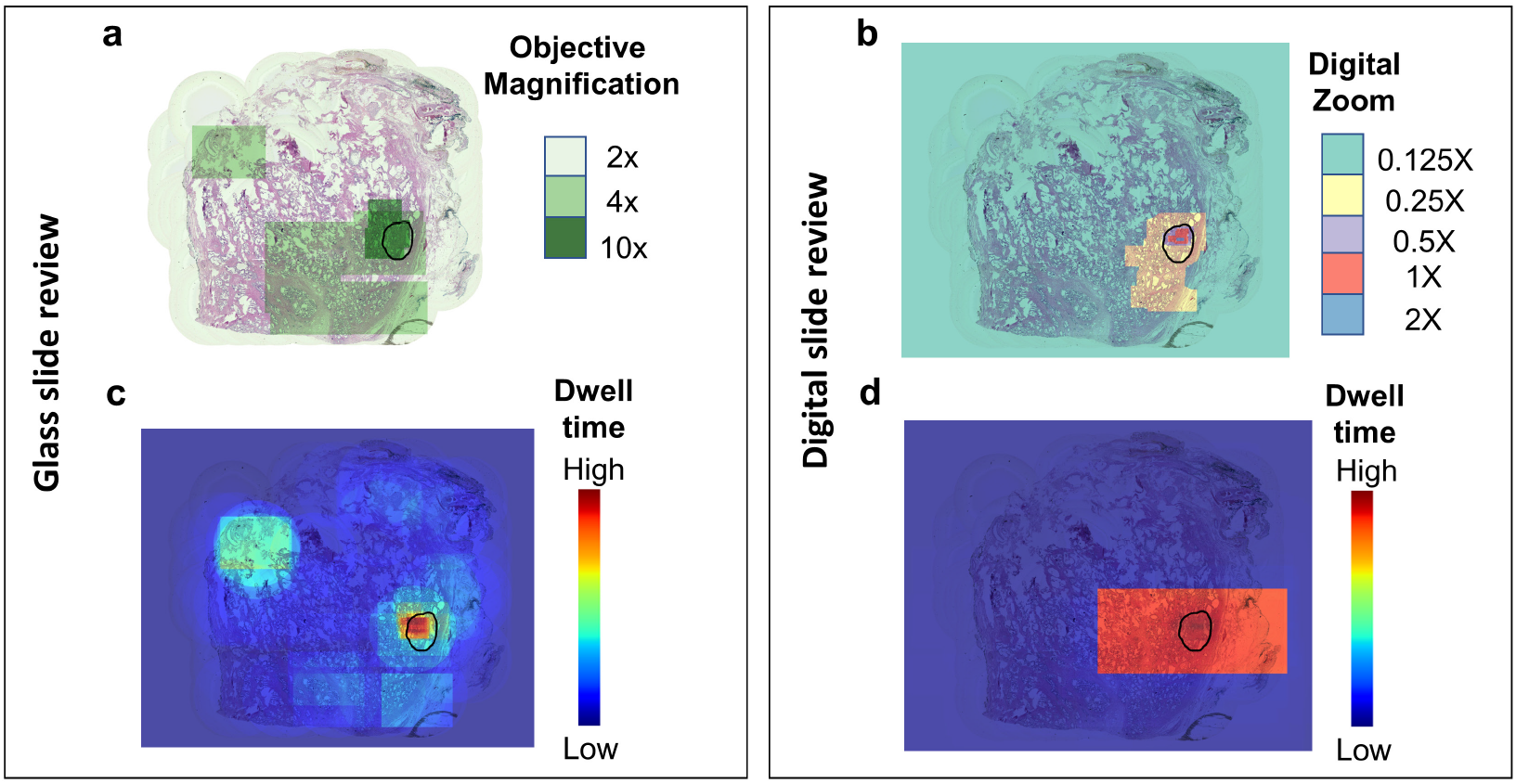
Digital review of PathCAM MRSIs after the glass slide diagnosis. When reviewing the glass slide during the PathCAM recording, the pathologist determined the Gleason grade to be 3+3. **a**, During the initial microscope recording, the pathologist observed the outlined tumor volume area at 10x magnification. **b**, The pathologist first reviewed the MRSI in HistomicsUI, where we captured the location of their viewport and zoom level over time. Then the pathologist annotated the tumor volume (black outlines). The highest digital zoom level was within the bounds of the tumor volume annotation. **c**, Multi-scale dwell-time map from the pathologist review of the glass slide at the microscope. **d**, Corresponding multi-scale dwell time map from pathologist review of the digital PathCAM MRSI.

**Supplementary Figure 7.**
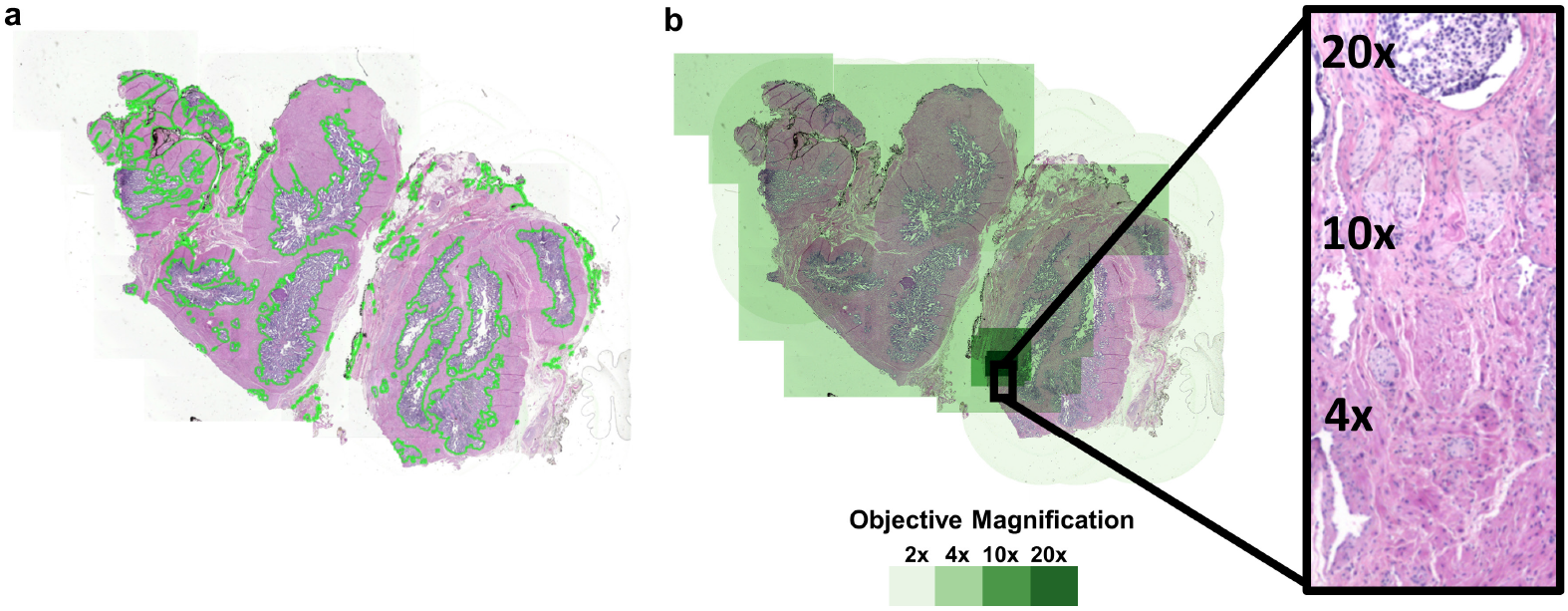
Semantic segmentation through HistomicsTK. **a**, We applied HistomicsTK’s color thresholding semantic segmentation to one of the multi-resolution slide iamges **b**,, which indicates areas of high cellularity (green outlines), to this MRSI to demonstrate the potential of these images to be used for additional computational analysis.

**Supplementary Figure 8.**
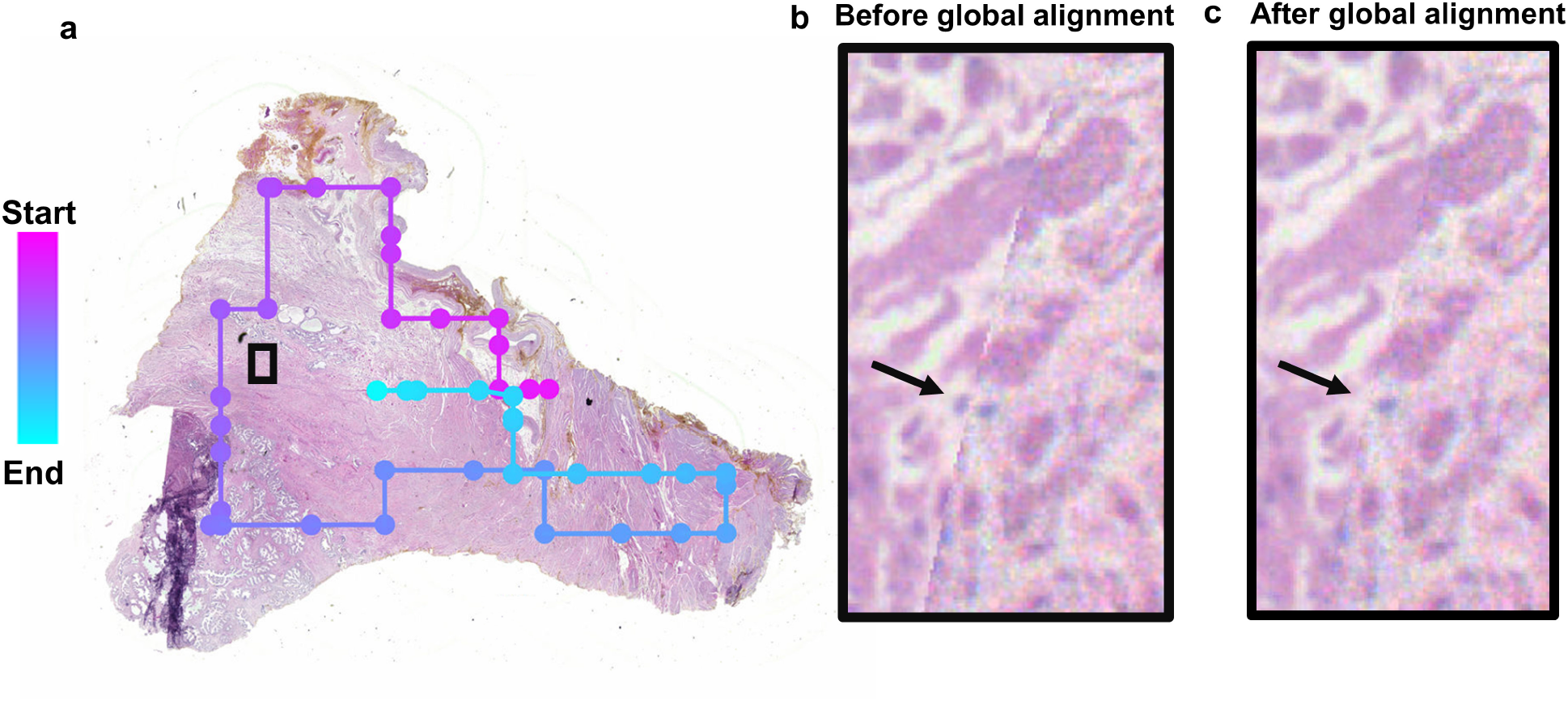
Global alignment for a 2X mosaic. **a**, The dots indicate the centroid of each registered frame along the search path. Dots in pink indicate the start of the mosaic sequence, and the color shifts to blue towards the end of the mosaic sequence. The black box shows an area of overlap due to a looping path. **b**, The black box region showing a misalignment (black arrow) before global alignment. **c**, The corrected frames after global alignment.

**Video 1.**
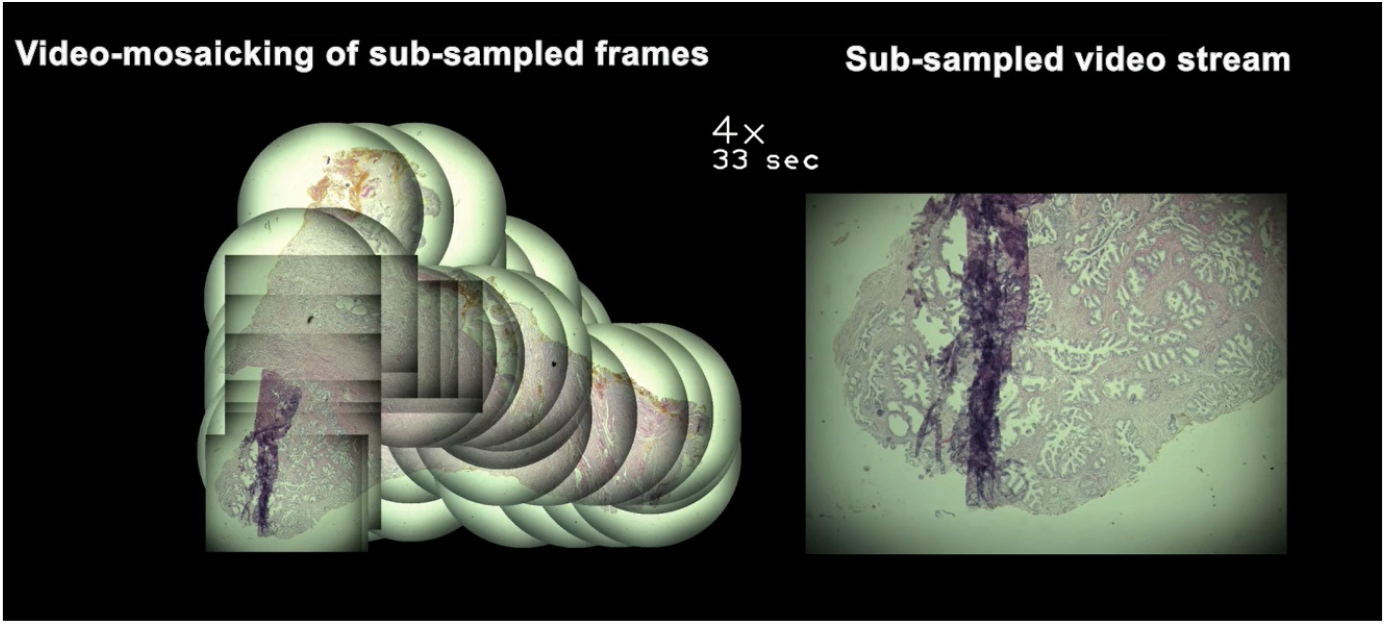

Demonstration of video-mosaicking of sub-sampled frames.

**Video 2.**
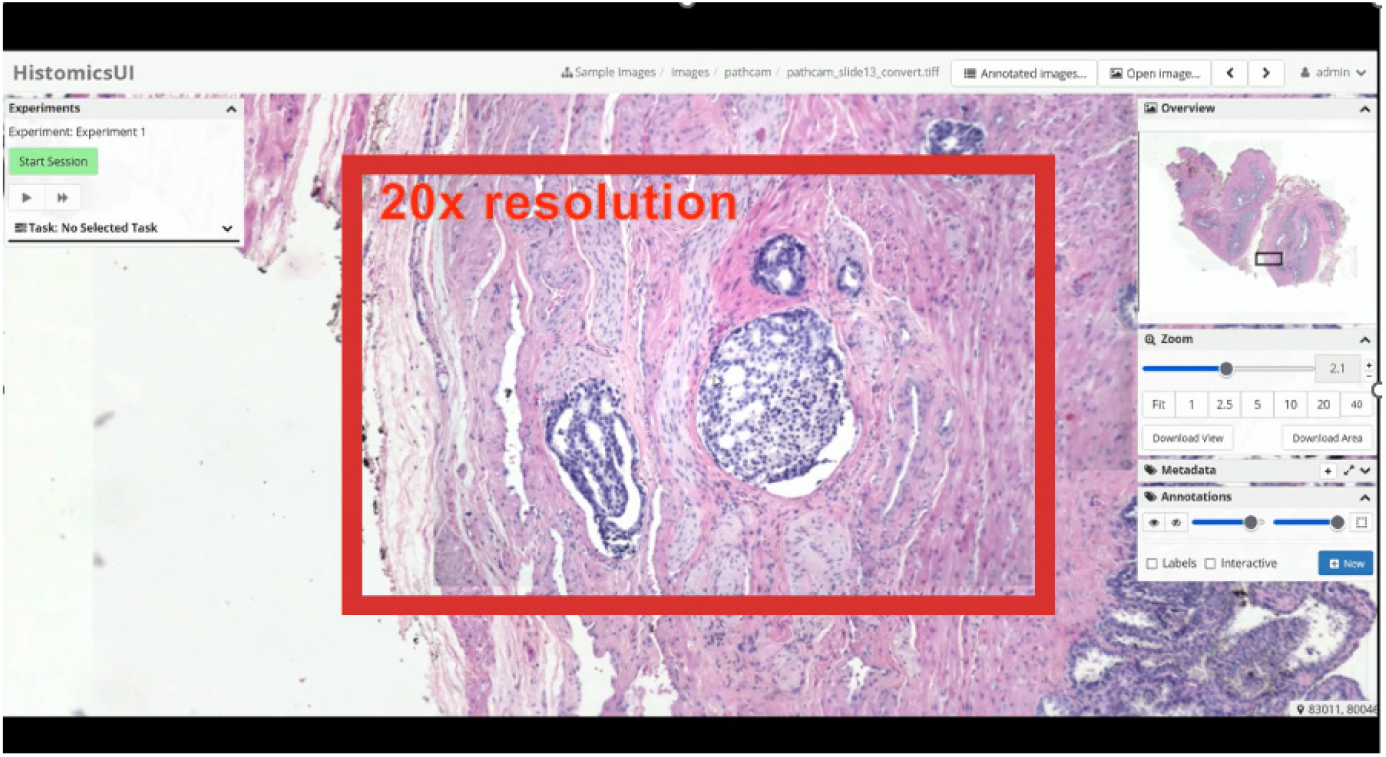

Demonstration of HistomicsUI interface with multi-resolution slide image.

**Video 3.**
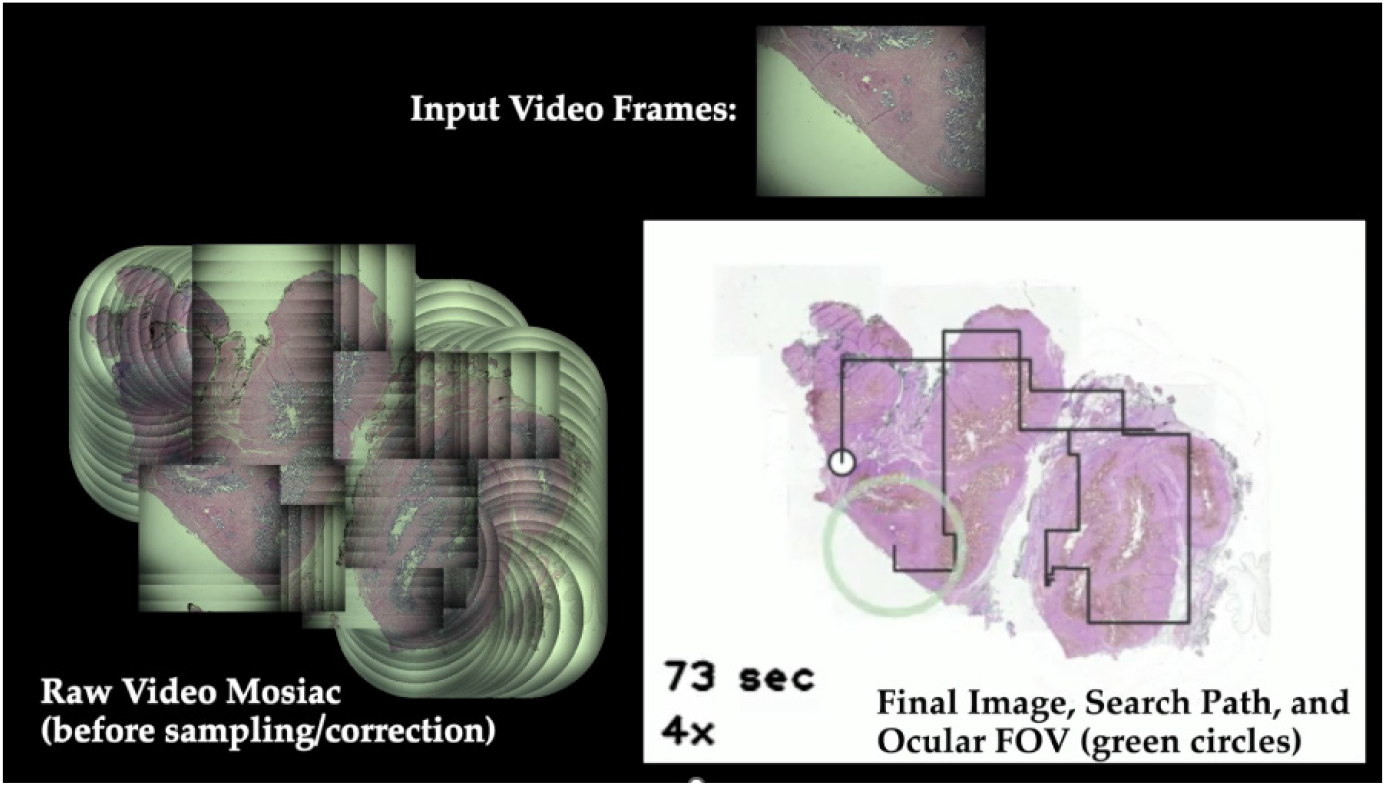

Demonstration of search path and ocular FOV over final multi-resolution slide image, compared with video and mosaic generation.

## Notes

### Competing Interest Statement

JQB is a shareholder and employee of Instapath, Inc., which did not financially or materially support this work. K.A., Z.H., B.S., C.W., M.C., and J.Q.B. are listed as inventors on a patent application related to this work.

